# Multifaceted suppression of staphylococcal virulence: *in vitro* study of a cellular status with reduced Agr activity and impaired biofilm

**DOI:** 10.1101/2025.11.11.687926

**Authors:** Nghi Bao Nguyen, Yuri Ushijima, Kazuya Morikawa, Le Thuy Thi Nguyen

## Abstract

Anti-quorum sensing (QS) therapy has been focused on to reduce the virulence of pathogenic bacteria. However, a series of studies have shown that suppression of the staphylococcal QS Agr system promotes biofilm formation. In this study, we found that both Agr activity and the biofilm of *Staphylococcus aureus* can be reduced by a sappanwood extract (SWe). The impaired biofilm in the presence of SWe exhibited reduced extracellular DNA (eDNA) and increased extracellular polysaccharides. SWe also reduced pigmentation. The RNA-seq analysis showed that SWe suppressed multiple key virulence regulators, including the Agr and SaeRS systems. SWe downregulated cell-wall-associated proteins and several Agr-regulated virulence factors such as hemolysins and phenol-soluble modulins (PSMs). The present study did not intend the clinical application of the SWe but exemplified that it is possible to achieve anti-Agr and anti-biofilm activities simultaneously, suggesting a new therapeutic concept to combat *S. aureus* infection.

## Introduction

*S. aureus* is a Gram-positive bacterium that asymptomatically inhabits the nasal cavities and skin surfaces of humans and animals, but it can cause diverse diseases ranging from food poisoning and superficial skin abscesses to severe ones such as pneumonia, meningitis, osteomyelitis, sepsis, and toxic shock syndrome (Humphreys, 2012). Methicillin-resistant *Staphylococcus aureus* (MRSA) is notorious for its resistance to *β*-lactams and poses a burden on a global scale (Guo et al., 2020; Murray et al., 2022). Hospital-associated MRSA (HA-MRSA) and Community-acquired MRSA (CA-MRSA) increased morbidity, mortality, and financial burdens on healthcare systems, necessitating comprehensive prevention and control measures (Crespo-Piazuelo & Lawlor, 2021; Guo et al., 2020; Murray et al., 2022). In 2019, it was estimated that more than 800,000 deaths were associated with antimicrobial resistance (AMR) globally by *S. aureus* (Guo et al., 2020). Moreover, livestock-associated MRSA (LA-MRSA) raises concerns about zoonotic transmission.

In addition to the antibiotic-resistance genes, *S. aureus* also forms the biofilm that protects microbes from host immune attacks and many antibiotics, making it fiendishly difficult to eradicate infections (Jacqueline & Caillon, 2014) (Paharik & Horswill, 2016). They are surrounded by a matrix of extracellular polysaccharides, extracellular and cell surface-associated proteins/adhesins, and extracellular DNA (eDNA) (Moormeier & Bayles, 2017). The main extracellular polysaccharide in *S. aureus* biofilm is poly-N-acetylglucosamine (PNAG), a.k.a. polysaccharide intercellular adhesin (PIA), which is synthesized by the enzymes encoded by *ica*ADBC operon (Nguyen et al., 2020). Although most *S. aureus* strains possess the *ica* genes, a subset does not depend on the PIA production to form biofilms, and the biofilm of *S. aureus* is divided into two groups: PIA-dependent biofilm and PIA-independent biofilm (McCarthy et al., 2015). PIA-dependent biofilm is mainly found in methicillin-sensitive *S. aureus* (MSSA), while MRSA prominently produces PIA-independent biofilm (Nguyen et al., 2020). The biofilm formation of *S. aureus* has been illustrated through the five development stages: attachment, multiplication, exodus, maturation, and dispersal (Moormeier & Bayles, 2017; Paharik & Horswill, 2016) (**Fig S1**).

The Agr system is the master regulator of virulence in *S. aureus* and is activated through a quorum-sensing/diffusion-sensing mechanism (**Fig S2**) (Le & Otto, 2015). In general, the Agr system turns on the expression of a series of secretory toxins/enzymes and turns off the expression of factors for surface attachment and biofilm formation. The Agr system is critical in invasive infection, and some inhibitors that target the Agr system are successful in pre-clinical trials (Ford et al., 2020; Tuchscherr & Otto, 2023; Vinodhini & Kavitha, 2024). On the other hand, Agr activity is thought to hinder chronic survival in the human body, and about half of the clinical isolates have no Agr activity due to mutations. Furthermore, we have reported that a small part of the Agr-negative strains can regain activity by a mechanism called phase variation, which may cause recurrent invasive infections (Gor et al., 2019). These indicate the difficulty in dealing with the staphylococcal infection strategies by inhibiting the Agr system alone.

A series of evidence established that the Agr system negatively regulates biofilm (Paharik & Horswill, 2016), and the suppression of Agr could lead to the promotion of biofilm **(Fig S1, S2**). At the early stage of biofilm formation, where the Agr is not activated, cell wall-associated proteins are expressed to facilitate cell attachment and local multiplication. In the dispersal stage, the Agr system plays a role in inducing exoproteins such as PSMs. It is also reported that the inactivation of Agr disables the release of cells from the *in vivo* biofilm in mouse model (Nishitani et al., 2015).

Through our ongoing efforts to find plant extracts applicable to the treatment of bacterial infections, the dichloromethane (DCM) extract of *Biancaea sappan* (formerly *Caesalpinia sappan* (Gagnon et al., 2016)) was found to have unexpected characteristics: it exhibited both anti-virulence and anti-biofilm activities in *S. aureus*. *Biancaea sappan*, commonly known as sappanwood or Indian redwood, is a tropical tree native to Southeast Asia. It has vibrant red heartwood that contains red pigments for dyeing textiles and producing red ink. The sappanwood tree has been used in traditional medicine to treat conditions such as inflammation and pain (Vij et al., 2023). In this study, we aimed to clarify the avirulent cellular status of *S. aureus,* where the Agr activity and biofilm are both impaired in response to the sappanwood extract.

## Results

### SWe impairs the biofilm

*S. aureus* SH1000 was selected as a representative strain to study the effect of the sappanwood extract (SWe). SH1000 is the derivative of the historically well-studied laboratory strain 8325-4 (Horsburgh et al., 2002). It is Agr positive and makes PIA-dependent biofilm (Horsburgh et al., 2002; Izano et al., 2008). The Minimal inhibition concentration (MIC) value of SWe on SH1000 was ∼500 µg/mL in MH broth and TSB (**Table S1**). In the biofilm assay conditions, where we inoculated a higher quantity of cells (10^6^ CFU per well) than in the MIC assay (10^5^ CFU per well), the CFU did not significantly change by 250 ug/mL SWe (95% compared with untreated control) (**Fig 1A**). Even at 500 µg/mL SWe, CFU was 80% of the untreated control, while the addition of MIC of chloramphenicol (62.5 µg/mL) reduced the CFU to an undetectable level. Thus, SWe is ineffective in its antibacterial activity in the biofilm assay conditions employed. Nevertheless, the biofilm of SH1000 was reduced by 25%, 35%, and 40% by 100, 250, and 500 µg/mL SWe (**Fig 1B**). The anti-biofilm effect of 250 µg/mL SWe was also observed in another medium, BHI + 3% NaCl, that is often employed for the analysis of biofilms (Chiba et al., 2022; O’Neill et al., 2007) (**Fig S3**). Thus, it is likely that SWe has anti-biofilm activity. Here, it must be noted that there are many reports where the growth inhibition was mistakenly regarded as the anti-biofilm activity. We therefore measured the CFU values of both biofilm and planktonic (washable) parts. If SWe has *bona fide* anti-biofilm effect, it would reduce biofilm CFU without reducing the CFU in the planktonic part. As expected, we could observe that the 250 µg/mL and 500 µg/mL SWe did not reduce the CFU value of planktonic cells while reducing that of biofilm (**Fig 1C**). The CFU in the planktonic part was increased in this lot of SWe, but it is not reproducible in different lots (data not shown). In summary, SWe interferes with SH1000 cells in biofilm development or maintenance without severe antimicrobial activity.

**Fig 1.**
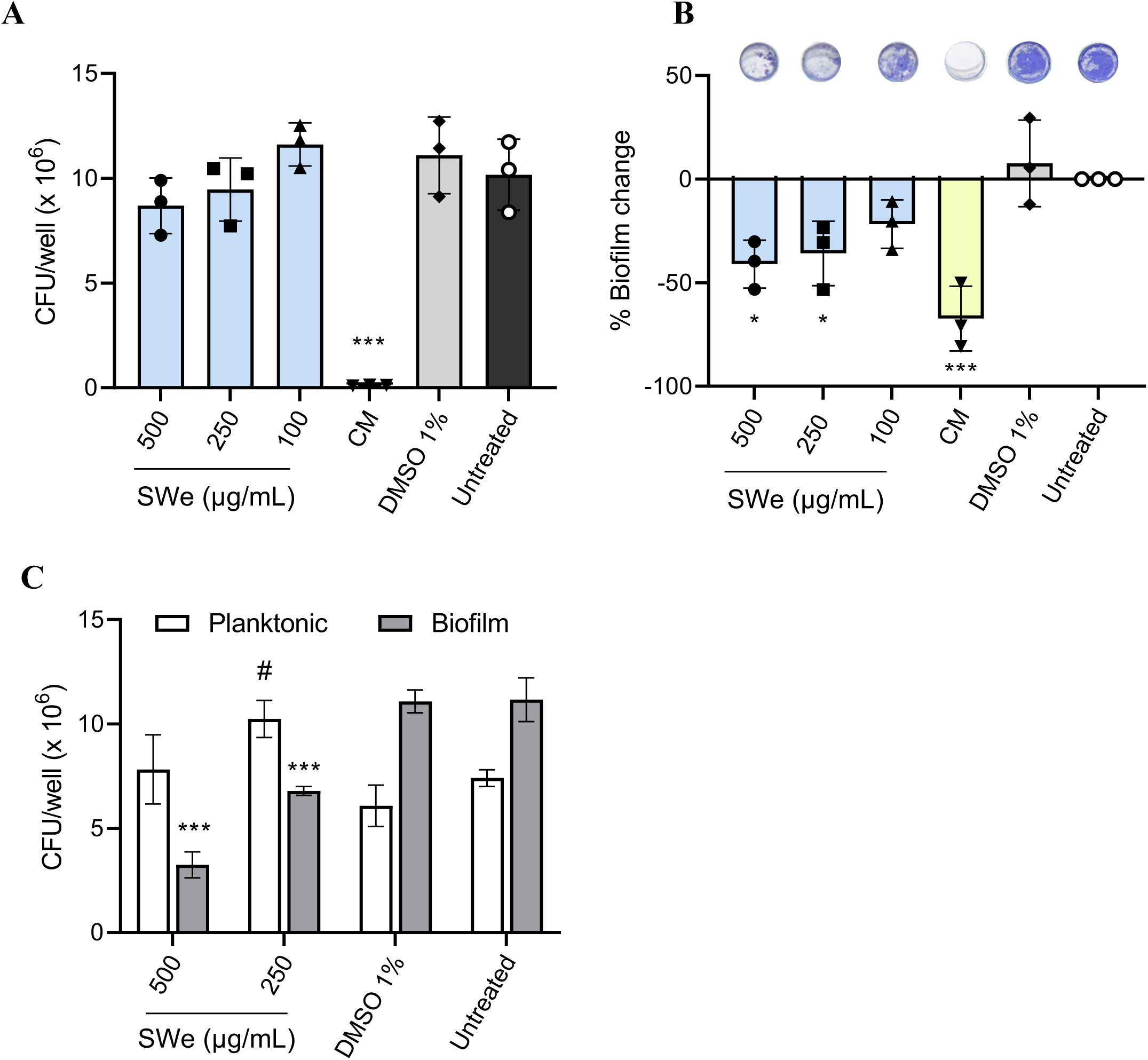
The antibiofilm activity of SWe on SH1000 after 24 hours of treatment. (A) Cell viability in total biomass (biofilm and planktonic cells) relative to untreated sample. The viable cells were counted as CFU. (B) Biofilm was quantified using crystal violet, and the values of percent change relative to the untreated group are shown. Representative images of stained biofilms are shown above the graph. (C) The viable cells were counted as CFU separately in planktonic (blank bard) and biofilm (filled bars) status. All data represent the mean values of 3 independent experiments; error bars indicate the SD. *: p≤ 0.05, **: p≤0.001, ***: p≤0.0001 compared to the untreated group, #: p≤ 0.05 planktonic cells compared with those in the untreated group. One-way ANOVA for (A, B) and Two-way ANOVA for (C) & Dunnett’s multiple comparison test with the untreated group. CM: 62.5 μg/mL chloramphenicol.

### SWe reduces eDNA in biofilm

Extracellular polysaccharides (EPS, also known as PIA), extracellular protein, and eDNA are the major constituents in *S. aureus* biofilm. These were quantified at different time points (3 h, 9 h, and 24 h) and were normalized by O.D._600_. Unexpectedly, PIA in the SWe treated group significantly increased at 9 h and 24 h (**Fig 2A**). The quantity of extracellular protein was not changed by SWe (**Fig 2B**). Instead, we observed a significant reduction in the eDNA amount in the SWe-supplemented biofilms throughout the tested time points (**Fig 2C**). The source of the eDNA is the genomic DNA released by the autolysis, and indeed, we found SWe reduces the autolysis (**Fig S4**). In addition, the biofilm degradation assay (**Fig S5**) also suggested the qualitative change in the biofilm.

**Fig 2.**
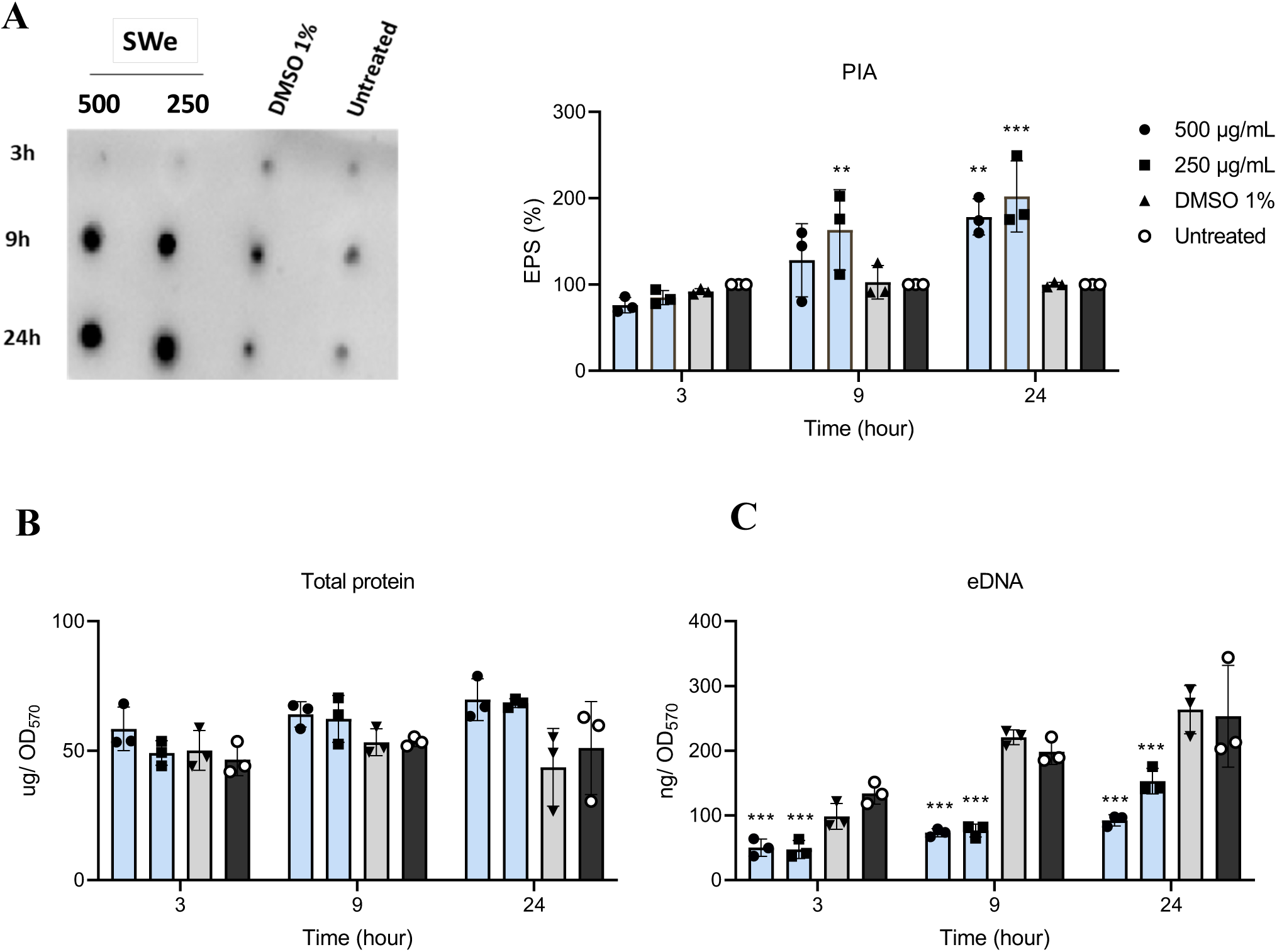
Quantification of ECM components in SH1000 biofilm treated with SWe at 3 h, 9 h, and 24 h. (A) PIA was detected by WGA staining (left). The signal intensity was quantified, and normalized by biofilm biomass (OD_570_), and the relative values compared with the untreated group are shown (right). (B) Extracellular protein (µg) normalized by OD_570_. (C) eDNA (ng) normalized by OD_570_. All data represent the average from 3 independent experiments; error bars indicate the SD. Two-way ANOVA & Dunnett’s multiple comparison test with the untreated group, *\*\**: *p≤0.001, \*\*\**: *p≤0.0001*.

### SWe suppresses the Agr activity

The effect of SWe on the Agr activity was tested using the reporter strain SH1000 P3*venus* in planktonic conditions (**Fig 3A, Fig S6**). The reporter plasmid expresses the fluorescence gene under the control of the *agr* P3 promoter. The control without SWe induced the fluorescence signal around 6h and then sustained a high level. On the other hand, the signal in the SWe-treated groups was suppressed. The suppression of fluorescence in the treated groups was not due to the delay or reduction of growth (**Fig 3B**). As observed under microscopy, the number of fluorescent cells was higher in the untreated group, and SWe reduced the fluorescent cells (**Fig S6**).

**Fig 3.**
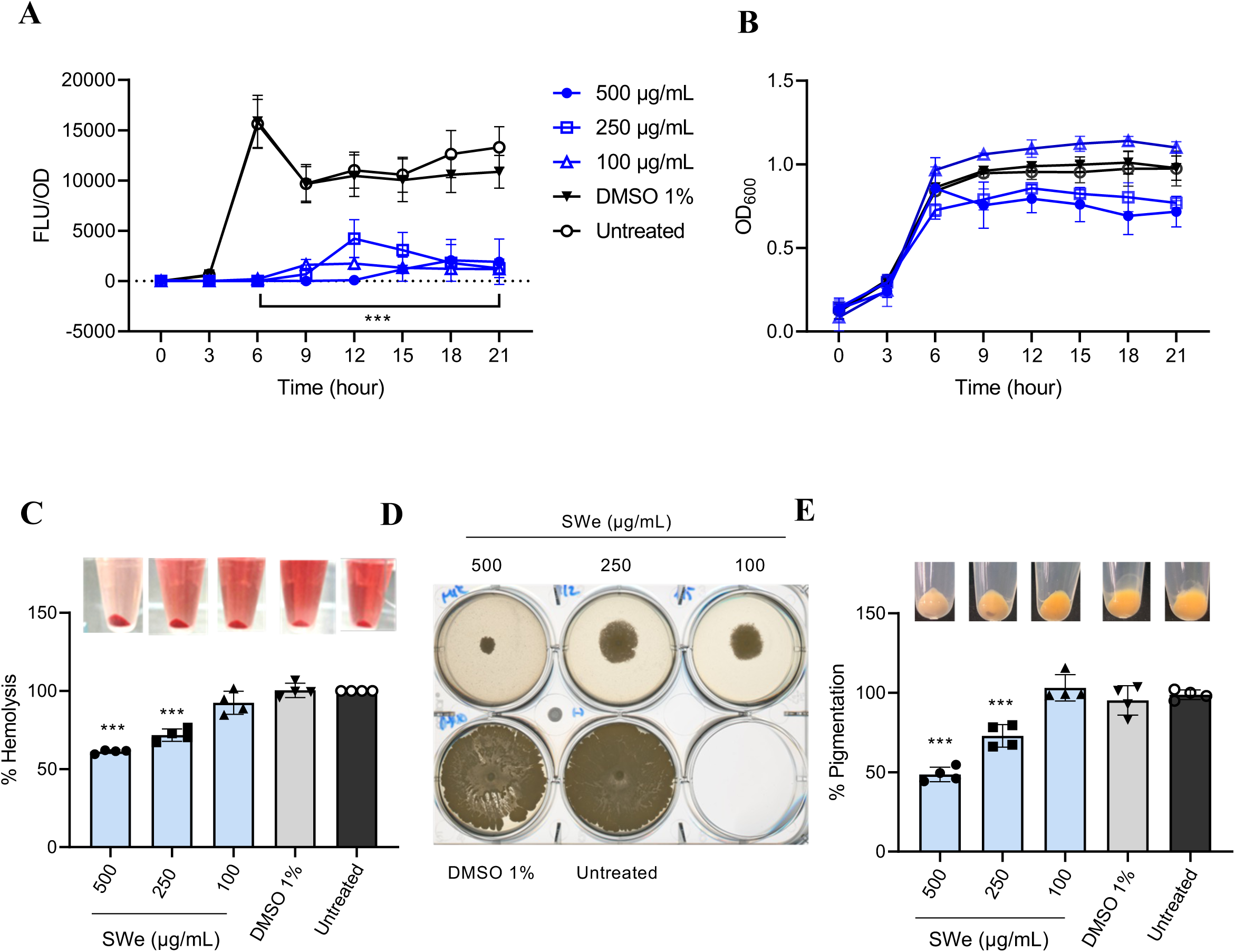
Inhibition of Agr system and virulence factors by SWe in SH1000 (18h). (A) The reporter signal was normalized by an OD_600_ value of SH1000 P3*venus*. (B) Growth curves of the reporter strain at different treatment concentrations. (C) Hemolysis activity of SH1000 after treating with SWe. (D) Colony spreading of SH1000. SWe was supplemented in the agar medium. (E) Pigmentation of SH1000. Photos of cell pellets after centrifugation are shown above the graph. All data represent the average of 3 or 4 independent experiments; error bars indicate the SD. Statistical significance (*\*\* p≤ 0.001, ***p ≤ 0.0001*) was observed for SWe treated groups (500, 250 and 100 μg/mL). One-way ANOVA for (C, E) and Two-way ANOVA for (A, B) & Dunnett’s multiple comparison test with the untreated group.

The anti-virulence activity of SWe against SH1000 was further tested in terms of hemolysis and colony spreading, both are regulated by the Agr system (**Fig S2**). The hemolytic activity of SH1000 was decreased by the addition of SWe to the growth medium, and the effect was dose-dependent (**Fig 3C**). Similarly, the spreading of SH1000 on soft agar, to which phenol soluble modulins (PSMs) mainly contribute (Kizaki et al., 2016), was suppressed by SWe (**Fig 3D**).

We also noticed that SWe reduces the yellow pigment production of SH1000 (**Fig 3E**). The carotenoids such as staphyloxanthin are virulence factors that support *S. aureus* survival under oxidative stress (Xue et al., 2019). The pigment in the treated SH1000 was significantly dropped by 50% and 35% at 500 and 250 µg/mL SWe, respectively. The pigmentation of *S. aureus* depends on the alternative sigma factor SigB (Kullik et al., 1998) and is not directly regulated by the Agr system, but some evidence shows the deletion of Agr reduces pigmentation (Marroquin et al., 2019; Tan et al., 2022).

### Cellular status in the presence of SWe revealed by RNA sequencing

So far, we have demonstrated that SWe can make *S. aureus* SH1000 into Agr-suppressed and biofilm-impaired status. The transcriptome was analyzed by RNA sequencing (RNA-seq) to gain additional insight into this interesting cellular status. Cells were statically grown as in biofilm assays, and all cells in each well (as a mixture of planktonic and biofilm cells) were submitted to the analysis. There were about 100-150 genes expressed differently (log_2_FC >1 and *p*-value < 0.05) in SWe conditions compared with controls at 3 h, 9 h, and 24 h (**Fig S7, Table S2-S4**). Downregulated and upregulated genes shared in at least 2 time points were 53 and 33, respectively (**Table 1-2**).

**Table 1.** Downregulated genes by SWe at two or more time points in 3 h, 9 h, and 24 h (*p* <0.05)

| Gene | Function | Gene expression |  |  |
| --- | --- | --- | --- | --- |
|  |  | 3h | 9h | 24h |
| efb | complement convertase inhibitor Efb | -7.91 | -3.07 | -6.05 |
| ecb | complement convertase inhibitor Ecb | -7.27 | -2.73 | -5.08 |
| scb | complement inhibitor SCIN-B | -6.55 | -1.99 | -4.99 |
| sbi | immunoglobulin-binding protein Sbi | -1.97 | -2.29 | -5.01 |
| eap | extracellular adherence protein Eap/Map | -4.13 | -3.75 | -2.47 |
| saeP | DM13 domain-containing protein | -3.92 | -2.43 | -4.29 |
| saeQ | DoxX family protein | -1.88 | -2.51 | -3.58 |
| sarY | SarA family transcriptional regulator | -3.33 | -1.27 | -2.49 |
| hlgC | bi-component gamma-hemolysin HlgCB subunit C | -1.77 | -3.26 | -3.25 |
| hyl | alpha-hemolysin | -3.17 | -2.25 | -5.48 |
| SAURSH1000_RS07670 | SAS049 family protein | -5.20 | -1.16 | -1.80 |
| SAURSH1000_RS05335 | beta-class phenol-soluble modulin | -4.80 | -1.09 | -2.95 |
| SAURSH1000_RS12605 | hypothetical protein | -5.57 | -3.42 | - |
| SAURSH1000_RS07170 | exodeoxyribonuclease VII small subunit | -5.31 | -5.68 | - |
| yhbY | ribosome assembly RNA-binding protein YhbY | -5.31 | -2.78 | - |
| SAURSH1000_RS12025 | hypothetical protein | -5.00 | -2.33 | - |
| SAURSH1000_RS03245 | hypothetical protein | -4.40 | -1.26 | - |
| SAURSH1000_RS07675 | CsbD family protein | -4.38 | -1.20 | - |
| SAURSH1000_RS03815 | DUF5067 domain-containing protein | -3.66 | -1.39 | - |
| SAURSH1000_RS00845 | isoprenylcysteine carboxyl methyltransferase family protein | -3.65 | -2.18 | - |
| SAURSH1000_RS08250 | hypothetical protein | -3.15 | -1.39 | - |
| SAURSH1000_RS00240 | phosphatidylinositol-specific phospholipase C | -2.81 | -1.95 | - |
| SAURSH1000_RS06945 | hypothetical protein | -2.76 | -2.26 | - |
| SAURSH1000_RS06090 | polymorphic toxin type 50 domain-containing protein | -2.56 | -1.94 | - |
| SAURSH1000_RS04915 | YkyA family protein | -2.30 | -1.33 | - |
| SAURSH1000_RS03810 | hypothetical protein | -2.01 | -1.76 | - |
| SAURSH1000_RS03270 | hypothetical protein | -1.87 | -1.16 | - |
| SAURSH1000_RS02080 | hypothetical protein | - | -2.66 | -5.45 |
| hlgB | bi-component gamma-hemolysin HlgAB/HlgCB subunit B | - | -2.60 | -2.24 |
| hlgA | bi-component gamma-hemolysin HlgAB subunit A | - | -1.89 | -4.07 |
| saeR | response regulator transcription factor SaeR | - | -1.87 | -2.12 |
| saeS | two-component system sensor histidine kinase SaeS | - | -1.53 | -2.14 |
| cntA | staphylopine-dependent metal ABC transporter substrate-binding protein CntA | - | -1.41 | -3.69 |
| SAURSH1000_RS08665 | DUF4888 domain-containing protein | - | -1.64 | -4.77 |
| SAURSH1000_RS02680 | long-chain fatty acid--CoA ligase | - | -2.55 | -3.33 |
| SAURSH1000_RS01930 | phenol-soluble modulin PSM-alpha-4 | -6.83 | - | -3.37 |
| SAURSH1000_RS01935 | phenol-soluble modulin PSM-alpha-2 | -6.75 | - | -3.53 |
| isdC | heme uptake protein IsdC | -6.29 | - | -2.29 |
| SAURSH1000_RS13275 | MAP domain-containing protein | -5.31 | - | -2.65 |
| SAURSH1000_RS05340 | beta-class phenol-soluble modulin | -5.24 | - | -4.24 |
| sarR | HTH-type transcriptional regulator SarR | -5.12 | - | -1.84 |
| SAURSH1000_RS13285 | delta-lysin family phenol-soluble modulin | -4.56 | - | -2.33 |
| sph | orotate phosphoribosyltransferase | -3.80 | - | -2.39 |
| sdrC | MSCRAMM family adhesin SdrC | -3.08 | - | -2.11 |
| SAURSH1000_RS08740 | tRNA-Ser | -2.91 | - | -3.50 |
| sdrD | MSCRAMM family adhesin SdrD | -2.70 | - | -2.01 |
| SAURSH1000_RS03975 | CsbD family protein | -2.43 | - | -2.35 |
| SAURSH1000_RS05245 | hypothetical protein | -2.17 | - | -3.24 |
| SAURSH1000_RS03135 | hypothetical protein | -2.10 | - | -1.75 |
| SAURSH1000_RS06035 | hypothetical protein | -1.84 | - | -2.58 |
| SAURSH1000_RS01685 | GlsB/YeaQ/YmgE family stress response membrane protein | -1.76 | - | -2.03 |
| SAURSH1000_RS01675 | PepSY domain-containing protein | -1.69 | - | -2.52 |
| SAURSH1000_RS03900 | histidine phosphatase family protein | -1.62 | - | -1.85 |

**Table 2.** Upregulated genes by SWe at two or more time points in 3 h, 9 h, and 24 h (*p* <0.05)

| Gene | Function | Log2FC |  |  |
| --- | --- | --- | --- | --- |
|  |  | 3h | 9h | 24h |
| SAURSH1000_RS06115 | ABC transporter ATP-binding protein | 7.40 | 4.85 | 2.39 |
| farE | fatty acid efflux MMPL transporter FarE | 7.23 | 4.47 | 5.88 |
| SAURSH1000_RS06120 | ABC transporter permease | 4.80 | 7.43 | 2.39 |
| SAURSH1000_RS12500 | hypothetical protein | 4.80 | 4.26 | 1.68 |
| SAURSH1000_RS12505 | DUF896 domain-containing protein | 3.93 | 4.53 | 1.68 |
| ptsG | glucose-specific PTS transporter subunit IIBC | 2.89 | 2.48 | 2.40 |
| SAURSH1000_RS06125 | sensor histidine kinase | 2.69 | 3.23 | 2.39 |
| ureB | urease subunit beta | 2.52 | 1.91 | 1.68 |
| ureA | urease subunit gamma | 2.33 | 2.09 | 1.68 |
| ureC | urease subunit alpha | 1.75 | 2.19 | 1.68 |
| ureD | urease accessory protein UreD | 1.73 | 2.21 | 1.68 |
| ureG | urease accessory protein UreG | 1.71 | 2.05 | 1.68 |
| SAURSH1000_RS06130 | response regulator | 2.39 | 3.91 | 2.39 |
| sceD | lytic transglycosylase SceD | 2.36 | 1.09 | 1.68 |
| SAURSH1000_RS01980 | sodium-dependent transporter | 3.19 | 1.20 | - |
| SAURSH1000_RS11325 | HlyD family efflux transporter periplasmic adaptor subunit | 3.14 | 1.50 | - |
| SAURSH1000_RS00800 | FMN-dependent NADH-azoreductase | 2.28 | 1.34 | - |
| SAURSH1000_RS02965 | tyrosine-type recombinase/integrase | 2.11 | 1.46 | - |
| SAURSH1000_RS06745 | PepSY domain-containing protein | 2.01 | 1.31 | - |
| SAURSH1000_RS13115 | SMP-30/gluconolactonase/LRE family protei | 1.69 | 1.03 | - |
| SAURSH1000_RS12525 | TetR/AcrR family transcriptional regulator | 1.68 | 1.39 | - |
| SAURSH1000_RS11460 | sucrose-specific PTS transporter subunit IIBC | 1.51 | 1.23 | - |
| SAURSH1000_RS11320 | DHA2 family efflux MFS transporter permease subunit | 3.36 | - | 2.18 |
| SAURSH1000_RS12275 | PTS fructose transporter subunit IIC | 2.32 | - | 1.72 |
| sdaAA | L-serine ammonia-lyase, iron-sulfur-dependent, subunit alpha | 2.15 | - | 2.05 |
| ureE | urease accessory protein UreE | - | 1.77 | 4.13 |
| ureF | urease accessory protein UreF | - | 1.61 | 4.06 |
| SAURSH1000_RS12330 | sterile alpha motif-like domain-containing protein | - | 1.59 | 2.77 |
| rpmB | 50S ribosomal protein L28 | - | 1.35 | 2.19 |
| SAURSH1000_RS06900 | HU family DNA-binding protein | - | 1.29 | 2.30 |
| fadA | thiolase family protein | - | 1.67 | 4.50 |
| fadB | 3-hydroxyacyl-CoA dehydrogenase/enoyl-CoA hydratase family protein | - | 1.26 | 4.22 |
| SAURSH1000_RS11850 | APC family permease | - | 1.06 | 1.68 |

The expression of *agr*BDCA operon was decreased at 3 h in RNA-seq data (**Fig 4A**). Consistently, the Agr-regulated virulence genes (*psma, psmb, hld, hla*) were suppressed even in the 24 h treatment (**Fig 4B**) and were confirmed by qPCR (**Fig 4C**).

**Fig 4.**
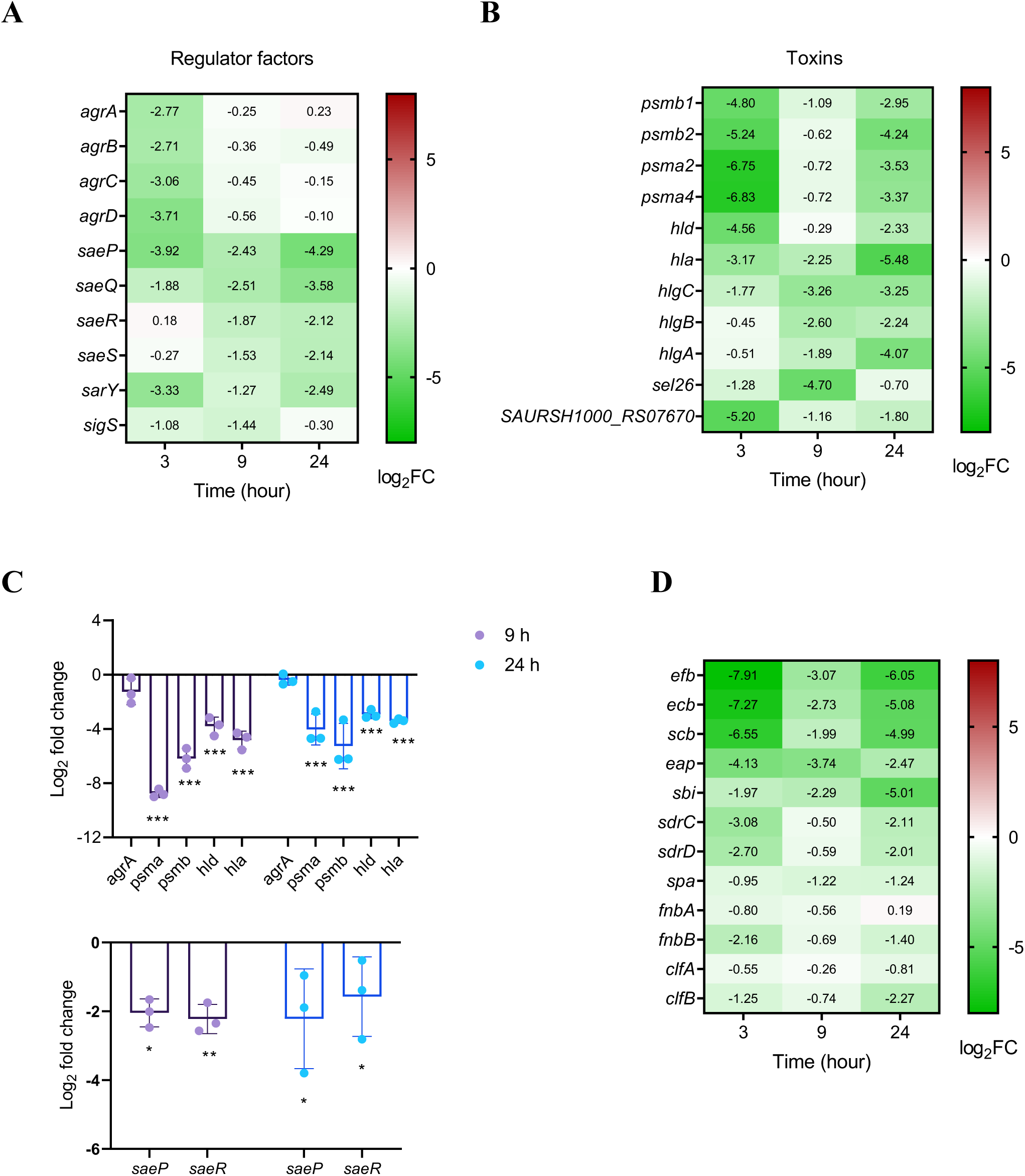
Differentially Expressed Genes (DEGs) in the SWe-treated cells: adhesins, exotoxins, and regulators. Log_2_ (Fold Change) values relative to untreated control at the same time point are shown. DEGs were grouped into regulator factors (**A**), exotoxins (**B**), and adhesins (**D**). (**C**) qPCR confirmation of the expression of selected genes at 9 h and 24 h. Error bars represent the SD; *\**: *p≤0.05, \*\**: *p≤0.001, \*\*\**: *p≤0.0001*. Fold change in expression levels was calculated by the 2^−ΔΔCT^ method. Statistical analysis was performed by unpaired t-test for the log_2_(fold change expression) values.

Increased PIA production (**Fig 2A**) and the decreased eDNA (**Fig 2B**) and autolytic activity (**Fig S4**) were not simply explained by our RNAseq analysis. In terms of PIA synthesis genes (*ica* operon), the expression was not increased in some genes and time points (**Fig S8**). Instead, we observed that the genes for the capsule biosynthesis pathway, which shares UDP-GlcNac as the starting material with the PIA biosynthesis pathway, were downregulated under treatment of SWe at 3 h(*capA* through *capP*) (**Fig S8**).

Many genes encoding adhesion proteins were downregulated in the SWe-treated sample (**Fig 4D**). This is one of the important findings in this study because adhesin proteins are known to be negatively regulated by the Agr system (Le & Otto, 2015). The evasion proteins extracellular fibrinogen-binding protein (Efb), extracellular complement-binding protein (Ecb), staphylococcal complement inhibitor B (Scb), staphylococcal binder of IgG (Sbi), and extracellular adherence protein (Eap) have been implicated in the *S. aureus* biofilm, possibly through positive charge–mediated interactions with eDNA and the cell wall (Dengler et al., 2015; Graf et al., 2019; Kavanaugh et al., 2019). Together with the low amount of eDNA in ECM (**Fig 2C**), the decrease in these adhesion proteins would affect the biofilm of *S. aureus* under treatment of SWe. Moreover, other MSCRAMMs involved in biofilm formation (FnBPA, FnnBPB, SdrC, SdrD, ClfA, ClfB, and SpA) tended to be lower in the SWe-treated sample. It is known that Efb, Sbi, Eap, FnBPA, FnBPB, and complement inhibitors are under the control of the two-component system SaeRS (Liu et al., 2016). Consistently, the expression of the SaePQRS was reduced by SWe (**Fig 4C, D**). SaeP is the bacterial cell surface protein required for the stability of biofilm by interaction with eDNA in ECM (Kavanaugh et al., 2019). Thus, the expression of some types of eDNA-binding proteins in ECM was reduced in SWe conditions, likely through the downregulation of the SaePQRS systems.

SWe also affected the expression of genes related to some metabolic pathways (**Fig S9**), and the intracellular level of reactive oxygen species (ROSs) (**Fig S10**), but their contribution to the anti-virulence and anti-biofilm activities is not known.

In summary, our transcriptome analysis confirmed the inhibitory effect of SWe on the Agr system. It also found the suppression of adhesin genes and the Sae system. Suppression of these virulence regulators is consistent with the SWe’s anti-biofilm and anti-virulence activities. Regarding the mechanism by which this ‘avirulent’ transcriptome status is achieved, it requires further extensive studies, including the identification of the active compounds. In the present study, we can only mention that the SWe’s major anti-biofilm effect is independent of the Agr system; SWe^lot2^ suppressed the biofilm in the absence of the Agr system (**Fig S11**).

### Effect of SWe on different types of *S. aureus* strains

We have shown that SWe affects key regulators and reduces biofilm and virulence of *S. aureus* SH1000, but the regulations of virulence expression (e.g., intactness of Agr system) and biofilm formation (e.g., PIA dependency) are diverse among strains. We tested the effect of SWe on two other strains, MW2 and ATCC25923. ATCC25923 is known for the medium biofilm formation and the lack of Agr function (Tan et al., 2015), in contrast, MW2 is a weak biofilm former with intact Agr function (Morales - laverde et al., 2022; Trotonda et al., 2008). Our assay confirmed these biofilm phenotypes (**Fig 5A**). SWe significantly reduced the biofilm of ATCC25923 but not of MW2 (**Fig 5A**). The anti-biofilm effect against ATCC25923 was not due to the growth inhibition (**Fig 5B**). Thus, SWe can disturb biofilm irrespective of the status of the Agr function.

**Fig 5.**
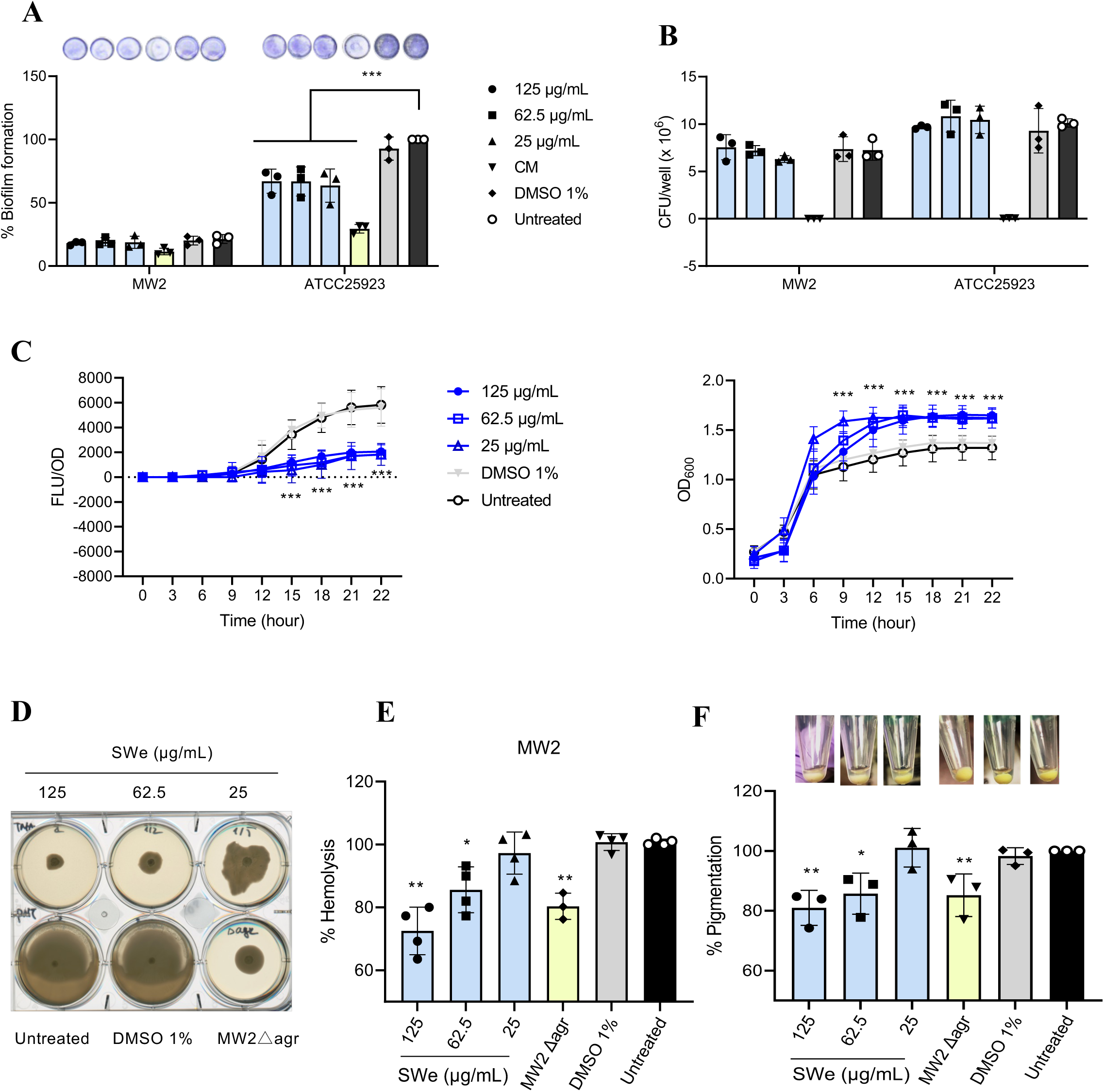
Effect of SWe on biofilm formation and virulence expression of MW2 and ATCC25923. Biofilm formation (A) and viability (B) of MW2 and ATCC25923 in the presence of SWe. The relative percentages to the untreated ATCC25923 biofilm is shown. (C) RNA-III reporter assay to test the anti-Agr activity of SWe on MW2. MW2 virulence phenotypes were also tested: (D) colony spreading, (E) hemolysis, and (F) pigmentation. All data represent the mean values of 3 independent experiments; error bars indicate the SD. Statistical significance (*: p≤ 0.05, **: p≤0.001, ***: p≤0.0001) was observed for SWe treated groups (125, 62.5 and 25 μg/mL). One-way ANOVA for (A, B, E, F) and two-way ANOVA for (C) & Dunnett’s multiple comparison test with the untreated group. CM: 62.5 μg/mL chloramphenicol.

In MW2, SWe suppressed the transcription of the RNA-III, as shown by the reporter assay (**Fig 5C**). Moreover, we also confirmed that SWe reduces hemolytic activity, colony spreading, and pigmentation (**Fig 5D-F**). These results suggest that SWe can reduce the virulence expression of MW2 without enhancing the biofilm formation. Overall, SWe can disturb the Agr system and impair biofilm in any type of strain tested. The effect of SWe on different strains is summarized in **Table 3**.

**Table 3.** Summary of the effect of SWe on *S. aureus* strains.

| Strain | Characteristics | Viability | Biofilm | Agr-dependent virulence |  |  | Pigmentation |
| --- | --- | --- | --- | --- | --- | --- | --- |
|  |  |  |  | RNA-III reporter | Hemolysis | Colony spreading |  |
| ATCC25923 | MSSA, *agr* type III (inactive), moderate biofilm | → | ↓ | not tested | not tested | not tested | not tested |
| SH1000 | MSSA, *agr* type I, moderate biofilm | → | ↓ | ↓ | ↓ | ↓ | ↓ |
| MW2 | MRSA, *agr* type III, weak biofilm | → | →<br>originally<br>weak | ↓ | ↓ | ↓ | ↓ |

## Discussion

SWe may not be immediately suitable for direct clinical use, as is the case with many natural plant products, but the significance of this study lies in providing the first example of a potential solution to overcome the limitations of current anti-quorum-sensing strategies against *S. aureus*. Inhibition of Agr can be achieved by a variety of inhibitors with different modes of action (Vinodhini & Kavitha, 2024). These anti-Agr activities are generally not through bactericidal activities or growth inhibition, which is an important feature for anti-virulence drugs that would not generate drug resistance. However, the inhibition of Agr results in increased biofilm. For example, staquorsin which is designed to interfere with the AgrA, promotes the biofilm formation of Newman (Mahdally et al., 2021). The extract of *Schinus terebinthifolia* fruit (450D-F5), and the purified compound (termed compound 1, oleanolic acid family) inhibit Agr and have a dose-dependent effect on biofilm but increase biofilm at low concentrations (Muhs et al., 2017) (Tang et al., 2020). Thus, no report has clearly shown the inhibition of both biofilm and Agr without affecting the growth. In this context, the present study first reported an example where the cells with suppressed Agr activity can sustain their planktonic lifestyle rather than shifting into the biofilm status, with the information about changes in the biofilm matrix and transcriptome underlying this interesting cellular status.

Regarding biofilm matrix compositions, SWe reduced eDNA (**Fig 2B**). The eDNA is the universal component in biofilm matrix not only in bacteria but also in fungi, with a negative charge in the structure to connect tightly to other molecular components in ECM (Campoccia et al., 2021; Liu et al., 2022; Shopova et al., 2013). Targeting eDNA for biofilm treatment has been focused on due to such a critical role (Campoccia et al., 2021). Previous studies have shown the relationship between eDNA and autolysis during biofilm formation (Biswas et al., 2006; Mann et al., 2009; Rice et al., 2007). We also found that SWe reduces autolytic activity (**Fig S4**), though the transcriptome analysis did not directly explain this. The SaeRS is known to affect the expression of *atl* encoding major autolysin (Mashruwala et al., 2017) and *nuc* encoding staphylococcal thermonuclease (Liu et al., 2016; Olson et al., 2013). However, the *atl* and *nuc* expressions were not changed by SWe. It might be valuable to note that autolysin can be inactivated by the local acidification at cell wall areas that are caused by the augmented respiration (Biswas et al., 2012). Cell wall metabolism and its regulator would also affect the cell lysis; for example, an alternative sigma factor SigB affects the cell wall thickness and changes the sensitivity to lysostaphin (cell wall degrading enzyme) (Morikawa et al., 2001). The present finding that SWe reduces the expression of *asp23* that is representative of the SigB regulon (**Fig S12**) may have implications in this aspect.

In addition to the decrease in eDNA, SWe increased PIA (**Fig 2A**), and altered the expression of genes encoding adhesin/cell surface proteins (**Fig 4A**). These qualitative changes in the biofilm matrix are likely the underlying mechanism in the SWe-dependent reduction in biofilm. Several virulence factors like Hla, Hld, and PSMs also contribute to the biofilm structure of *S. aureus* (Graf et al., 2019; Schwartz et al., 2012; Tayeb-Fligelman et al., 2017). SWe affected the expression of these factors in Agr-positive strains, SH1000 and MW2 (**Fig 5**).

The present study found that SWe negatively affects the SaeRS system. The SaeRS system has been known as the critical regulator of biofilm (Mashruwala et al., 2017; Moormeier & Bayles, 2017; Mrak et al., 2012) (**Fig S1**). Some reports showed that SaeRS targeting compounds reduce biofilm in *S. aureus*. An example that inhibits SaeR is Fenoprofen, a nonsteroidal anti-inflammatory drug (trade name Nalfon) (Jiang et al., 2023). Fenoprofen can restore the walking of mice with implant-associated osteomyelitis. The Fenoprofen-treated biofilm structure was loosened and porous due to the reduction of eDNA and proteins. The PIA production was unchanged, and its antibiofilm activity was not recorded in the PIA-dependent biofilm type, unlikely in our SWe case.

The carotenoid pigments are among the virulence factors to cope with oxidative stress (Clauditz et al., 2006). The expression of genes encoding CrtMN carotenoid synthesis enzymes is controlled by the alternative sigma factor SigB (Xue et al., 2019). Our RNA-seq analysis detected the SWe-dependent reduction in expression of *asp23* (representative of SigB regulon) at 3h (**Table S2, Fig S12**), but not in *crtMN*. Several studies have shown the involvement of some signaling and metabolisms in pigmentation, e.g., Agr system, respiration, and metabolic pathways (Lan et al., 2010) (Schurig-Briccio et al., 2020) (Marroquin et al., 2019). SWe-dependent reduction in pigmentation is likely a part of the alteration in cellular metabolism and signaling, where ROS and the Agr system are involved. Alternatively, some molecules in SWe might be the inhibitors for the carotenoid synthesis enzymes, likely as suggested mechanism for the farnesol inhibition of the carotenoid synthesis (Vila et al., 2019).

The methanol and ethanol extract of sappanwood had bactericidal activity but no anti-QS activity (data not shown), unlikely to the DCM solvent used in this study (SWe). This suggests that the anti-QS compound(s) in the DCM extract has less polarity. Sappanwood contains a variety of compounds, including xanthone, coumarin, phenolic compounds, chalcones, flavones, homo isoflavonoids, and brazilin (Vij et al., 2023). Peng et al. show the antibiofilm effect of brazilin by upregulating *icaR* and *agrA* expression (Peng et al., 2018), which is different from the effect of SWe in this study. Sappanwood extracted in DCM may also contain fatty acids (Ngernnak et al., 2018), which can show antibiofilm activity (Alenazy, 2023; Lee et al., 2017). Bioactive compound(s) in SWe are to be identified in our ongoing studies to understand the molecular input(s) that transform *S. aureus* avirulent. In addition, exploring the other extracts or compounds that can suppress both Agr and biofilm, followed by the analysis of the cellular status, would be an important comparative study to unravel the general cellular response that can comprehensively maintain *S. aureus* avirulent.

## Materials and methods

### Extraction from *B. sappan*

*B. sappan* wood was purchased from a Traditional medicine pharmacy in Ho Chi Minh City. It was finely ground into a powder. Ten grams of the powder were soaked in 100 mL of Dichloromethane (DCM) (270997, Merck, Germany) solvent and allowed to extract for 24 hours. The extract was then filtered by Whatman filter paper to collect the liquid phase. This DCM extraction process was repeated twice, combining the resulting extracts. The solvent was subsequently evaporated to yield the DCM extract. The obtained extract was dissolved in 100% dimethyl sulfoxide (DMSO) (4987481246928, FUJIFIRM Wako, Japan) to achieve 50 mg/mL, which served as the stock solution for subsequent experiments (designated as SWe). We used this concentration as the approximate reference; SWe still contains DMSO-insoluble materials, including the wood powder. Insoluble materials were removed by 0.22µm membrane filter (for SWe lot number 1) (Minisart-NY25 17845 ACK, Sartorius, Germany), or centrifugation at 10,000rpm for 3min (for SWe lot number 2). The extract was checked for its antibiofilm activity. The ones with confirmed antibiofilm and anti-Agr activities were selected and used in this study (lot numbers 1 and 2). SWe-lot1 is used for all main experiments, and SWe-lot2 was used in the Supplementary Figures S5 and S11 since SWe-lot1 was not available anymore.

### Bacterial strains and culture conditions

The strains used in this study are listed in **Table 4**. Strains SH1000, MW2 and ATCC25923 are from the laboratory stock collection. *S. aureus* SH1000 served as the model strain to explore the anti-biofilm and anti-virulence properties of SWe. The deletion mutant of the *agr* locus (SH1000Δagr) was constructed by using pMAD-tet-Δagr (Gor et al., 2019). Cells were inoculated in Tryptic Soy Broth (TSB) (00382902571612, BD, USA) or Mueller Hinton Broth (MHB) (00382902757306, BD, USA) from −80°C glycerol stocks and grown at 37°C with shaking at 180 rpm overnight. Chloramphenicol (12.5 μg/mL) (036-10571, FUJIFILM Wako, Japan) was supplemented to TSB for the overnight cultures of reporter strains, SH1000-RNA III-gfp and MW2-RNA III-gfp.

**Table 4.**
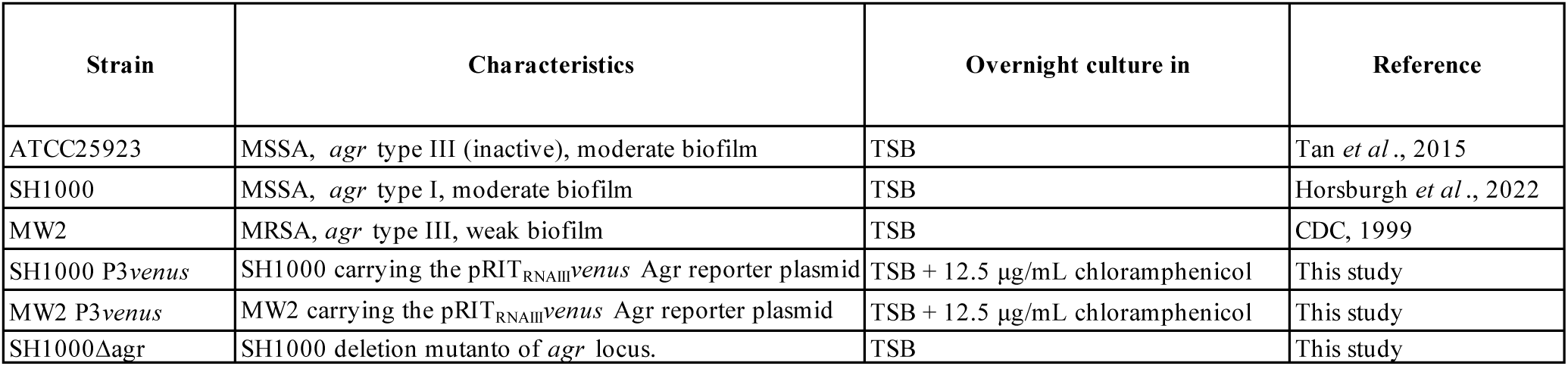
List of strains and overnight culture condition.

| Strain | Characteristics | Overnight culture in | Reference |
| --- | --- | --- | --- |
| ATCC25923 | MSSA, *agr* type III (inactive), moderate biofilm | TSB | Tan *et al.*, 2015 |
| SH1000 | MSSA, *agr* type I, moderate biofilm | TSB | Horsburgh *et al.*, 2022 |
| MW2 | MRSA, *agr* type III, weak biofilm | TSB | CDC, 1999 |
| SH1000 P3*venus* | SH1000 carrying the pRIT<sub>RNAIII</sub>*venus* Agr reporter plasmid | TSB + 12.5 µg/mL chloramphenicol | This study |
| MW2 P3*venus* | MW2 carrying the pRIT<sub>RNAIII</sub>*venus* Agr reporter plasmid | TSB + 12.5 µg/mL chloramphenicol | This study |
| SH1000Δ*agr* | SH1000 deletion mutanto of *agr* locus. | TSB | This study |

### MIC

The MIC value was determined following the protocol of the CLSI standard (“CLSI M26-A - Methods for Determining Bactericidal Activity of Antimicrobial Agents; Approved Guideline,”). Bacteria cultured overnight in MHB were diluted to a 10^6^ CFU/mL concentration. A volume of 100 μL of the bacterial suspension (10^5^ CFU) was distributed into each well of a 96-well microtiter plate. Successive two-fold dilutions of the extract SWe, ranging from 2000 μg/mL (4% volume of SWe) to 0 μg/mL, were prepared in MHB. Subsequently, 100 μL of each diluted extract was added to the corresponding wells. DMSO, at a concentration of 4%, served as the solvent control and was diluted similarly to the extract. Bacterial cultures without any additives were also tested. The microtiter plate was incubated at 37°C for 24 hours. The MIC value was determined as the lowest concentration of the extract that inhibited visible bacterial growth.

### Biofilm assay and viability test (96-well plates)

The biofilm assay protocol was adapted from Gomes et al. (Gomes et al., 2019). *S. aureus* was cultivated in 96-well plates (260860, Thermo, Denmark) (**Figs 1, 2, 7)**. Each well contained 100 μL of TSB and 10^6^ CFU of *S. aureus*. Appropriate concentrations of SWe were supplemented. Control groups included TSB alone, TSB with DMSO solvent controls (1% DMSO is equivalent to SWe at 500 µg/mL prepared from 50 mg/mL stock), and TSB with 62.5 µg/mL of Chloramphenicol. Each condition was tested in triplicate in each independent experiment. The plates were incubated at 37°C for 24 hours. After incubation, planktonic bacteria were removed, and the wells were washed twice with phosphate-buffered saline (PBS). Biofilms were fixed by adding 100 μL of 90% ethanol to each well. After 2 min incubation, the ethanol was removed and the plate was air-dried. The biofilms were stained with 0.1% crystal violet solution for 5 min, after which the stain was removed, and the wells were air-dried and washed with PBS. The absorbed crystal violet dye in biofilms was solubilized in 100 μL of 90% ethanol for 15 min on the horizontal shaker. The harvested crystal violet was measured as absorbance at 570 nm.

To evaluate the effect of SWe on SH1000 growth (**Fig 1C)**, both the supernatant and biofilm were collected after 24 hours of incubation. The biofilm structures were disrupted by pipetting. CFU was counted by using TSB agar (TSA) plates.

### Biofilm assay and viability test (optimized for 24-well plates)

*S. aureus* was cultivated in 24-well plates (3526, Corning, USA) (**Supplementary Figures S5, S11)**. Each well contained 300 μL of TSB and 3 × 10^6^ CFU of *S. aureus*. Planktonic bacteria were removed, and wells were washed twice with phosphate-buffered saline (PBS). To evaluate the effect of SWe on cell growth (**Fig S11)**, both the supernatant and biofilm were collected after 24 hours of incubation.

For CV staining (**Fig S5**), protocol was optimized for the 24-well plate system. Washed biofilms were first air-dried and fixed by adding 500 μL of 80% ethanol to each well. After 2 min incubation, ethanol was diluted in a stepwise manner (80%, 40%, and 5%). The biofilms were stained with 0.1% solution of crystal violet (031-07331, FUJIFILM Wako, Japan) for 5 min. The stained biofilm was washed once with water, and the plate image was scanned. The absorbed crystal violet dye in biofilms was solubilized in 600 μL of 90% ethanol with 1.5% HCl. The harvested crystal violet was measured as absorbance.

### Quantification of ECM components

The extraction and quantification of the ECM were conducted following a modified version of the method described by Chiba et al. (Chiba et al., 2015). Biofilms were collected from 96-well plates and dissolved in 1.5 M NaCl, followed by a 5-min incubation at room temperature. The samples were then centrifuged at 5,000 rpm for 10 min to separate the supernatants. These supernatants were subsequently analyzed for PIA content, eDNA, and protein concentration.

PIA was quantified using Wheat Germ Agglutinin (WGA), a carbohydrate-binding lectin with a high affinity for sialic acid and N-acetylglucosamine. This lectin stains yeast buds as well as the cell surfaces of gram-positive bacteria and animal cells (Wang et al., 2016). A 5 μL aliquot of each sample was dot-blotted onto a nitrocellulose membrane. The membrane was blocked with 1% Bovine Serum Albumin (BSA) and incubated with HRP-WGA reagent (29073, Biotium, USA) at a concentration of 130 ng/mL in 1% BSA solution. Detection of PIA was performed using an Enhanced Chemiluminescence (ECL) solution (1705060, Bio-Rad, USA) and visualized with a FUSION chemiluminescence, fluorescence, and gel imaging system (Vilber Lourmat, Germany). The signal of EPS was processed by ImageJ software, then the EPS (%) was calculated: (signal/area)_treated_ /(signal/area)_untreated_ × 100.

eDNA was quantified using the Quant-iT™ PicoGreen® dsDNA Reagent and Kit (P11496, Thermo Fisher, USA) according to the manufacturer’s instructions. Protein concentration was measured using the DC™ Protein Assay Kit I (5000111, Bio-Rad, USA).

### Bacterial lysis assay

The overnight culture of SH1000 was diluted 100-fold in TSB. SWe (final concentration of 250 μg/mL) was added to 5 mL of the prepared bacterial suspension in a 15 mL tube. Tubes were incubated at 37°C with shaking at 180 rpm for 3 hours, after which the bacterial cells were harvested by centrifugation at 10,000 rpm for 2 min at room temperature. The resulting pellets were washed twice with PBS. Subsequently, the pellets were suspended in 1 mL PBS and adjusted to OD_600_ of 0.2–0.3. A volume of 100 μL of each adjusted suspension was added to a 96-well plate as triplicate for each sample. The OD_578_ was monitored every 15 min for a total of 3 hours with shaking at 180 rpm.

### DNase I, Protease K, and metaperiodate treatment of biofilm

Biofilm stability against DNase I (LS002138, Worthington, USA), Protease K(166-28913, FUJIFILM Wako, Japan), and sodium metaperiodate (1.06597, Merk, Germany) was tested as previously described (Seidl et al., 2008) (Sugimoto et al., 2018) with some modifications. The cells were inoculated in a 24-well plate (300 µl TSB medium per well) and incubated at 37°C. After 24 hours, DNase I (140U/mL, final conc.), Protease K (100 µg/mL), or sodium metaperiodate (10 or 50 mM) (Seidl et al., 2008) was directly added to each well, followed by an additional 2 hours of incubation at 37°C.

In separate wells, DNase I was added to the culture upon inoculation, and the biofilm formation in the presence of DNase I was observed (**Fig S5**, right column).

Following the treatment, biofilm was quantified by CV as described above (optimized method for 24-well plates).

### Effect of extract treatment on hemolytic activity

An overnight culture of *S. aureus* SH1000 was diluted to approximately 10⁷ CFU/mL in TSB supplemented with or without SWe and the solvent control 1% DMSO. Cultures were incubated at 37 °C with shaking at 180 rpm for 16 –18 h. After incubation, cells were pelleted by centrifugation at 10,000 rpm for 3 min, and the supernatants were collected. A 250 µL aliquot of each supernatant was mixed with 250 µL of a 2% sheep red blood cell suspension prepared in PBS 1X. The mixtures were incubated for 1 h at 4 °C, followed by centrifugation at 500 rpm for 5 min to pellet intact red blood cells. Hemolytic activity was quantified by measuring the absorbance of the supernatants at 540 nm.

### Effect of SWe on the colony spreading

Colony spreading was performed as described by Kaito et al. (Kaito & Sekimizu, 2007). SH1000 was cultured at 37°C in TSB for 16–18 hours with shaking at 180 rpm. Subsequently, 2 μL of the bacterial culture was spotted onto 0.24%-agar TSA plates supplemented with SWe at MIC, 1/2 MIC, and 1/5 MIC. Control groups included no SWe or 1% DMSO (solvent control). All plates were incubated at 37°C for 16–18 hours.

### Carotenoids measurement

The carotenoid pigments were extracted and quantified as described (Morikawa et al., 2001). An overnight culture of SH1000 was diluted to 10^7^ CFU/mL in TSB supplemented with SWe. Negative controls consisted of untreated bacteria and those treated with 1% DMSO. Cultures were incubated at 37°C for 16–18 hours with shaking at 180 rpm. Following incubation, cells were harvested by centrifugation at 10,000 rpm for 1 min, and washed twice with sterile water at room temperature. The cells were suspended in 500 μL of methanol and incubated at 55°C for 5 min. After centrifugation, the supernatants were collected. The methanol extraction was repeated once more. The absorbance of the combined supernatants was measured at 465 nm using a plate reader with a 100 μL sample. The percentage of pigmentation was calculated relative to the untreated control group.

### Fluorescence reporter assay of Agr activity

Overnight cultures of SH1000 P3*venus* and MW2 P3*venus* reporter strains were diluted to 10^7^ CFU/mL in TSB supplemented with SWe. Controls are TSB or TSB supplemented with 1% DMSO. A 100 μL aliquot of each bacterial suspension was added to 96-well plates in triplicate. The OD at 590 nm and fluorescence (FL) signal (485 – 520 nm) were measured using an Infinite F Nano+ plate reader (Tecan, Switzerland) every 15 min for 25 hours with an orbital shaking amplitude set of 4.

### RNA extraction for qPCR and RNA-seq

Cells were grown under static conditions as previously described. At 3, 9, and 24 hours, the cell suspensions, including both planktonic and biofilm cells, were harvested in new tubes and centrifuged at 10,000 rpm for 3 min at 4°C. The cell pellets were washed twice with suspension buffer (50 mM Tris, pH 8.0, 1 mM EDTA, 50 mM NaCl) and resuspended in 200 μL of the same buffer. Five microliters of 10 mg/mL lysostaphin (120-06611, FUJIFIRM Wako, Japan) was added to the suspension and incubated at 37°C for 10 min, followed by the addition of 1 mL ice-cold TRIzol reagent (93289-100ML, Sigma, USA). The mixture was incubated on ice for 5 min and was centrifuged at 13,000 rpm for 20 min at 4°C. The upper aqueous phase was collected, and 200 μL of cold chloroform was added. The mixture was vortexed at maximum speed for 3 min, left for 3 min to facilitate phase separation, and centrifuged at 13,000 rpm for 15 min at 4°C. The upper phase was collected and mixed with an equal volume of cold isopropanol. The mixture was incubated at −20°C for 20 min, followed by centrifugation at 15,000 rpm for 20 min at 4°C to pellet the RNA. The RNA pellet was washed twice with cold 70% ethanol, air-dried for 10 min, and dissolved in 30 μL of DEPC-treated water. The extracted RNA was quantified by NanoDrop™ OneC UV-Vis Spectrophotometer (Thermo Fisher, USA).

### qPCR

Following RNA extraction, the samples were treated with DNase and reverse transcribed into cDNA using the PrimeScript RT reagent Kit with gDNA Eraser (RR047A, Takara, Japan) according to the manufacturer’s instructions. TB Green Premix Ex Taq II Fast qPCR kit (RR830A, Takara, Japan) was used for qPCR. Three separate samples were each tested in triplicate. The relative fold change in gene expression was determined using the 2^−ΔΔCt^ method using 16S RNA as the normalization reference. The primers used for detecting gene expression are listed in Table 5.

**Table 5.**
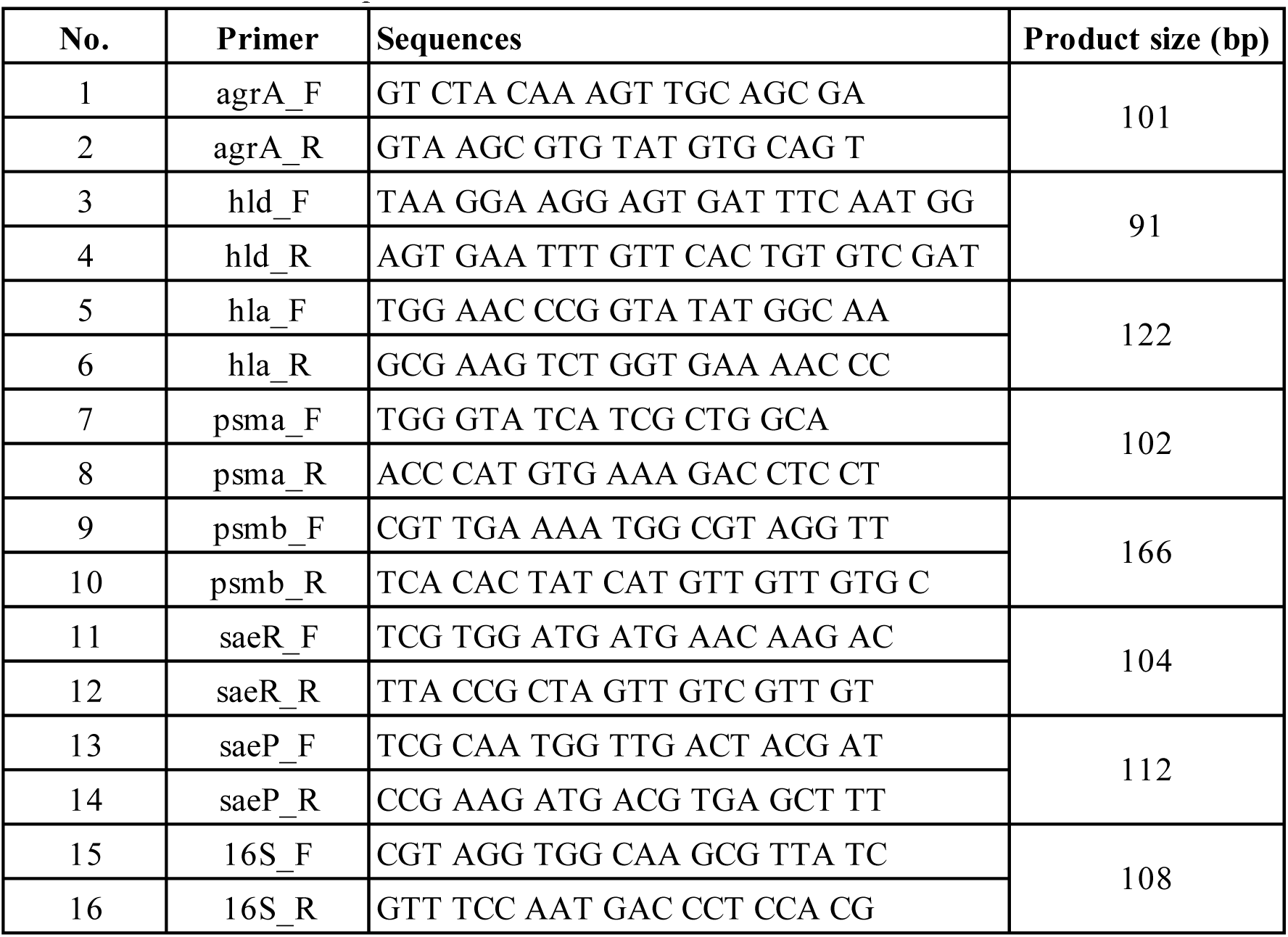
Primer list for qPCR.

### RNA-seq

RNA-seq was done by Rhelixa company (Japan). rRNA was removed from total RNA using a RiboZero Plus rRNA Depletion kit (Epicentre, USA). Transcripts were fragmented and used as a template to generate a strand-specific cDNA library using the NEBNext Ultra II Directional RNA Library Prep kit (NEB, USA). The sequencing was conducted on the Illumina NovaSeq6000 system with 150bp paired-end reads. RNA-seq raw reads were trimmed by Trimmomatic software and aligned to chromosomes of *S. aureus* SH1000 (NZ_CP059180.1) by HISAT2 (Kim et al., 2018). Mapped reads were counted using ‘featureCounts’ (Liao et al., 2014). Differentially expressed genes between the SWe-treated sample and untreated sample were analyzed with edgeR (Chen, 2014) with the threshold *p*<0.05 (Benjamini-Hochberg method) and |log_2_foldchange| > 1.

### NO and ROS detection

The biofilm cells at 9 h and 24 h were harvested and centrifuged at 15,000 rpm for 2 min at room temperature. The cells were washed once and resuspended in 1 mL of PBS. For plate reader analysis, the cell pellet was adjusted to OD_600_ of approximately 0.2 and divided into two tubes, each containing 500 μL, to separately detect ROS and NO. To detect intracellular ROS, CellROX™ Orange Reagent (C10443, Thermo Fisher, USA) was added to a final concentration of 5 μM and incubated for 30 min at 37°C in the dark. For NO detection, 5 μM DAF-FM diacetate reagent (16296, Cosmo Bio, Japan) was added and incubated for 1 hour under the same condition. After incubation, cells were pelleted and washed twice with PBS. The cells were resuspended in 500 μL of PBS, and ROS and NO levels were measured at 545–565 nm and 485–520 nm, respectively, using Infinite® M Nano (Tecan) plate reader.

For microscopy analysis, DAF-FM diacetate reagent was added to the cell suspension at a final concentration of 5 μM and incubated for 30 min at 37°C in the dark. Subsequently, CellROX™ Orange Reagent was added to the same tube at the final concentration of 5 μM and further incubated for 30 min. Cells were washed and resuspended in 1 mL of PBS. Two microliters of the suspension were loaded onto a glass slide and observed under a fluorescent microscope (BZ-X710, Keyence).

### Statistical analysis

All assays were repeated by at least 3 independent experiments. Statistical analysis was performed with GraphPad Prism (version 8.0) software (La Jolla, CA). Statistical methods are shown in figure captions.

## Acknowledgement

We sincerely thank Dr. Pham Thi Kim Tram of the Department of Medical Biotechnology, Biotechnology Center of Ho Chi Minh City, for generously providing laboratory facilities that enabled us to perform some experimental work in Vietnam during April 2025.

## References

Alenazy, R. (2023). Antimicrobial Activities and Biofilm Inhibition Properties of *Trigonella foenumgraecum* Methanol Extracts against Multidrug-Resistant *Staphylococcus aureus* and *Escherichia coli*. Life 2023, Vol. 13, Page 703, 13(3), 703–703. 10.3390/LIFE13030703

Biswas, R., Martinez, R. E., Göhring, N., Schlag, M., Josten, M., Xia, G., Hegler, F., Gekeler, C., Gleske, A. K., Götz, F., Sahl, H. G., Kappler, A., & Peschel, A. (2012). Proton-binding capacity of *Staphylococcus aureus* wall teichoic acid and its role in controlling autolysin activity. PloS one, 7(7). 10.1371/JOURNAL.PONE.0041415

Biswas, R., Voggu, L., Simon, U. K., Hentschel, P., Thumm, G., & Götz, F. (2006). Activity of the major staphylococcal autolysin Atl. FEMS microbiology letters, 259(2), 260–268. 10.1111/J.1574-6968.2006.00281.X

Campoccia, D., Montanaro, L., & Arciola, C. R. (2021). Extracellular DNA (eDNA). A Major Ubiquitous Element of the Bacterial Biofilm Architecture. International Journal of Molecular Sciences, 22(16). 10.3390/IJMS22169100

CDC. (1999). From the Centers for Disease Control and Prevention. Four pediatric deaths from community-acquired methicillin-resistant *Staphylococcus aureus*--Minnesota and North Dakota, 1997-1999. JAMA, 282(12), 1123–1125. https://www.ncbi.nlm.nih.gov/pubmed/10501104

Chen, Y., Lun, A. T., & Smyth, G. K. (2014). Differential expression analysis of complex RNA-seq experiments using edgeR.. In Statistical analysis of next generation sequencing data (1st ed., pp. 51–74). Springer International Publishing : Imprint: Springer,. 10.1007/978-3-319-07212-8_3

Chiba, A., Seki, M., Suzuki, Y., Kinjo, Y., Mizunoe, Y., & Sugimoto, S. (2022). *Staphylococcus aureus* utilizes environmental RNA as a building material in specific polysaccharide-dependent biofilms. NPJ Biofilms Microbiomes, 8(1), 17. 10.1038/s41522-022-00278-z

Chiba, A., Sugimoto, S., Sato, F., Hori, S., & Mizunoe, Y. (2015). A refined technique for extraction of extracellular matrices from bacterial biofilms and its applicability. Microbial biotechnology, 8(3), 392–403. 10.1111/1751-7915.12155

Clauditz, A., Resch, A., Wieland, K. P., Peschel, A., & Götz, F. (2006). Staphyloxanthin plays a role in the fitness of *Staphylococcus aureus* and its ability to cope with oxidative stress. Infection and Immunity, 74(8), 4950–4953. 10.1128/IAI.00204-06/ASSET/0B500F87-F8F6-43EC-8C7E-02D1C33F109E/ASSETS/GRAPHIC/ZII0080661110005.JPEG

CLSI M26-A - Methods for Determining Bactericidal Activity of Antimicrobial Agents; Approved Guideline. In.

Crespo-Piazuelo, D., & Lawlor, P. G. (2021). Livestock-associated methicillin-resistant *Staphylococcus aureus* (LA-MRSA) prevalence in humans in close contact with animals and measures to reduce on-farm colonisation. Irish Veterinary Journal 2021 74:1, 74(1), 1–12. 10.1186/S13620-021-00200-7

Dengler, V., Foulston, L., DeFrancesco, A. S., & Losick, R. (2015). An Electrostatic Net Model for the Role of Extracellular DNA in Biofilm Formation by *Staphylococcus aureus*. Journal of Bacteriology, 197(24), 3779–3779. 10.1128/JB.00726-15

Ford, C. A., Hurford, I. M., & Cassat, J. E. (2020). Antivirulence Strategies for the Treatment of *Staphylococcus aureus* Infections: A Mini Review. Frontiers in Microbiology, 11, 14–14. 10.3389/FMICB.2020.632706

Gagnon, E., Bruneau, A., Hughes, C. E., de Queiroz, L. P., & Lewis, G. P. (2016). A new generic system for the pantropical *Caesalpinia* group (*Leguminosae*). PhytoKeys(71), 1–160. 10.3897/phytokeys.71.9203

Gomes, F., Martins, N., Ferreira, I. C. F. R., & Henriques, M. (2019). Anti-biofilm activity of hydromethanolic plant extracts against *Staphylococcus aureus* isolates from bovine mastitis. Heliyon, 5(5). 10.1016/J.HELIYON.2019.E01728

Gor, V., Takemura, A. J., Nishitani, M., Higashide, M., Romero, V. M., Ohniwa, R. L., & Morikawa, K. (2019). Finding of Agr Phase Variants in *Staphylococcus aureus*. mBio, 10(4). 10.1128/MBIO.00796-19

Graf, A. C., Leonard, A., Schäuble, M., Rieckmann, L. M., Hoyer, J., Maass, S., Lalk, M., Becher, D., Pané-Farré, J., & Riedel, K. (2019). Virulence Factors Produced by *Staphylococcus aureus* Biofilms Have a Moonlighting Function Contributing to Biofilm Integrity. Molecular & Cellular Proteomics : MCP, 18(6), 1036–1036. 10.1074/MCP.RA118.001120

Guo, Y., Song, G., Sun, M., Wang, J., & Wang, Y. (2020). Prevalence and Therapies of Antibiotic-Resistance in *Staphylococcus aureus*. Frontiers in cellular and infection microbiology, 10. 10.3389/FCIMB.2020.00107

Horsburgh, M. J., Aish, J. L., White, I. J., Shaw, L., Lithgow, J. K., & Foster, S. J. (2002). σB Modulates Virulence Determinant Expression and Stress Resistance: Characterization of a Functional *rsb*U Strain Derived from *Staphylococcus aureus* 8325-4. Journal of Bacteriology, 184(19), 5457–5457. 10.1128/JB.184.19.5457-5467.2002

Humphreys, H. (2012). *Staphylococcus aureus:* the enduring pathogen in surgery. The surgeon : journal of the Royal Colleges of Surgeons of Edinburgh and Ireland, 10(6), 357–360. 10.1016/J.SURGE.2012.05.003

Izano, E. A., Amarante, M. A., Kher, W. B., & Kaplan, J. B. (2008). Differential Roles of Poly-N-Acetylglucosamine Surface Polysaccharide and Extracellular DNA in *Staphylococcus aureus* and *Staphylococcus epidermidis* Biofilms. Applied and Environmental Microbiology, 74(2), 470–470. 10.1128/AEM.02073-07

Jacqueline, C., & Caillon, J. (2014). Impact of bacterial biofilm on the treatment of prosthetic joint infections. The Journal of antimicrobial chemotherapy, 69 Suppl 1(SUPPL1). 10.1093/JAC/DKU254

Jiang, F., Chen, Y., Yu, J., Zhang, F., Liu, Q., He, L., Musha, H., Du, J., Wang, B., Han, P., Chen, X., Tang, J., Li, M., & Shen, H. (2023). Repurposed Fenoprofen Targeting SaeR Attenuates *Staphylococcus aureus* Virulence in Implant-Associated Infections. ACS Central Science, 9(7), 1354–1373. 10.1021/ACSCENTSCI.3C00499/SUPPL_FILE/OC3C00499_SI_002.XLSX

Kaito, C., & Sekimizu, K. (2007). Colony Spreading in *Staphylococcus aureus*. Journal of Bacteriology, 189(6), 2553–2553. 10.1128/JB.01635-06

Kavanaugh, J. S., Flack, C. E., Lister, J., Ricker, E. B., Ibberson, C. B., Jenul, C., Moormeier, D. E., Delmain, E. A., Bayles, K. W., & Horswill, A. R. (2019). Identification of extracellular DNA-binding proteins in the biofilm matrix. mBio, 10(3). 10.1128/MBIO.01137-19/SUPPL_FILE/MBIO.01137-19-ST002.DOCX

Kim, T., Seo, H. D., Hennighausen, L., Lee, D., & Kang, K. (2018). Octopus-toolkit: a workflow to automate mining of public epigenomic and transcriptomic next-generation sequencing data. Nucleic Acids Res, 46(9), e53. 10.1093/nar/gky083

Kizaki, H., Omae, Y., Tabuchi, F., Saito, Y., Sekimizu, K., & Kaito, C. (2016). Cell-Surface Phenol Soluble Modulins Regulate *Staphylococcus aureus* Colony Spreading. PLoS ONE, 11(10). 10.1371/JOURNAL.PONE.0164523

Kullik, I., Giachino, P., & Fuchs, T. (1998). Deletion of the alternative sigma factor σ(B) in *Staphylococcus aureus* reveals its function as a global regulator of virulence genes. Journal of Bacteriology, 180(18), 4814–4820. 10.1128/JB.180.18.4814-4820.1998/ASSET/328A9147-5F90-4DC9-8702-FFFDAD4877F9/ASSETS/GRAPHIC/JB1880515005.JPEG

Lan, L., Cheng, A., Dunman, P. M., Missiakas, D., & He, C. (2010). Golden pigment production and virulence gene expression are affected by metabolisms in *Staphylococcus aureus*. Journal of bacteriology, 192(12), 3068–3077. 10.1128/JB.00928-09

Le, K. Y., & Otto, M. (2015). Quorum-sensing regulation in staphylococci-an overview. Frontiers in Microbiology, 6(OCT), 167362–167362. 10.3389/FMICB.2015.01174/BIBTEX

Lee, J. H., Kim, Y. G., Park, J. G., & Lee, J. (2017). Supercritical fluid extracts of *Moringa oleifera* and their unsaturated fatty acid components inhibit biofilm formation by *Staphylococcus aureus*. Food Control, 80, 74–82. 10.1016/J.FOODCONT.2017.04.035

Liao, Y., Smyth, G. K., & Shi, W. (2014). featureCounts: an efficient general purpose program for assigning sequence reads to genomic features. Bioinformatics, 30(7), 923–930. 10.1093/bioinformatics/btt656

Liu, Q., Yeo, W. S., & Bae, T. (2016). The SaeRS Two-Component System of *Staphylococcus aureus*. Genes, 7(10). 10.3390/GENES7100081

Liu, S., Le Mauff, F., Sheppard, D. C., & Zhang, S. (2022). Filamentous fungal biofilms: Conserved and unique aspects of extracellular matrix composition, mechanisms of drug resistance and regulatory networks in *Aspergillus fumigatus*. npj Biofilms and Microbiomes 2022 8:1, 8(1), 1–8. 10.1038/s41522-022-00347-3

Mahdally, N. H., George, R. F., Kashef, M. T., Al-Ghobashy, M., Murad, F. E., & Attia, A. S. (2021). Staquorsin: A Novel *Staphylococcus aureus* Agr-Mediated Quorum Sensing Inhibitor Impairing Virulence i*n vivo* Without Notable Resistance Development. Front Microbiol, 12, 700494. 10.3389/fmicb.2021.700494

Mann, E. E., Rice, K. C., Boles, B. R., Endres, J. L., Ranjit, D., Chandramohan, L., Tsang, L. H., Smeltzer, M. S., Horswill, A. R., & Bayles, K. W. (2009). Modulation of eDNA Release and Degradation Affects *Staphylococcus aureus* Biofilm Maturation. PLOS ONE, 4(6), e5822–e5822. 10.1371/JOURNAL.PONE.0005822

Marroquin, S., Gimza, B., Tomlinson, B., Stein, M., Frey, A., Keogh, R. A., Zapf, R., Todd, D. A., Cech, N. B., Carroll, R. K., & Shaw, L. N. (2019). MroQ is a novel Abi-domain protein that influences virulence gene expression in *Staphylococcus aureus* via modulation of *agr* activity. Infection and Immunity, 87(5). 10.1128/IAI.00002-19/SUPPL_FILE/IAI.00002-19-S0001.PDF

Mashruwala, A. A., Gries, C. M., Scherr, T. D., Kielian, T., & Boyd, J. M. (2017). SaeRS Is Responsive to Cellular Respiratory Status and Regulates Fermentative Biofilm Formation in *Staphylococcus aureus*. Infection and immunity, 85(8). 10.1128/IAI.00157-17

McCarthy, H., Rudkin, J. K., Black, N. S., Gallagher, L., O’Neill, E., & O’Gara, J. P. (2015). Methicillin resistance and the biofilm phenotype in *Staphylococcus aureus*. Front Cell Infect Microbiol, 5, 1. 10.3389/fcimb.2015.00001

Moormeier, D. E., & Bayles, K. W. (2017). *Staphylococcus aureus* biofilm: a complex developmental organism. Molecular microbiology, 104(3), 365–376. 10.1111/MMI.13634

Moormeier, D. E., Bose, J. L., Horswill, A. R., & Bayles, K. W. (2014). Temporal and stochastic control of *Staphylococcus aureus* biofilm development. mBio, 5(5). 10.1128/MBIO.01341-14

Morales-laverde, L., Echeverz, M., Trobos, M., Solano, C., & Lasa, I. (2022). Experimental Polymorphism Survey in Intergenic Regions of the *ica*ADBCR Locus in *Staphylococcus aureus* Isolates from Periprosthetic Joint Infections. Microorganisms, 10(3). 10.3390/MICROORGANISMS10030600

Morikawa, K., Maruyama, A., Inose, Y., Higashide, M., Hayashi, H., & Ohta, T. (2001). Overexpression of sigma factor, sigma(B), urges *Staphylococcus aureus* to thicken the cell wall and to resist beta-lactams. Biochem Biophys Res Commun, 288(2), 385–389. 10.1006/bbrc.2001.5774

Mrak, L. N., Zielinska, A. K., Beenken, K. E., Mrak, I. N., Atwood, D. N., Griffin, L. M., Lee, C. Y., & Smeltzer, M. S. (2012). *sae*RS and *sar*A act synergistically to repress protease production and promote biofilm formation in *Staphylococcus aureus*. PloS one, 7(6). 10.1371/JOURNAL.PONE.0038453

Muhs, A., Lyles, J. T., Parlet, C. P., Nelson, K., Kavanaugh, J. S., Horswill, A. R., & Quave, C. L. (2017). Virulence Inhibitors from Brazilian Peppertree Block Quorum Sensing and Abate Dermonecrosis in Skin Infection Models. Scientific Reports 2017 7:1, 7(1), 1–15. 10.1038/srep42275

Murray, C. J. L., Ikuta, K. S., Sharara, F., Swetschinski, L., Robles Aguilar, G., Gray, A., Han, C., Bisignano, C., Rao, P., Wool, E., Johnson, S. C., Browne, A. J., Chipeta, M. G., Fell, F., Hackett, S., Haines-Woodhouse, G., Kashef Hamadani, B. H., Kumaran, E. A. P., McManigal, B., … Naghavi, M. (2022). Global burden of bacterial antimicrobial resistance in 2019: a systematic analysis. Lancet (London, England), 399(10325), 629–655. 10.1016/S0140-6736(21)02724-0

Ngernnak, C., Panyajai, P., Anuchapreeda, S., Wongkham, W., & Saiai, A. (2018). PHYTOCHEMICAL AND CYTOTOXIC INVESTIGATIONS OF THE HEARTWOOD OF *CAESALPINIA SAPPAN* LINN. Asian Journal of Pharmaceutical and Clinical Research, 11(2), 336–339. 10.22159/AJPCR.2018.V11I2.22903

Nguyen, H. T. T., Nguyen, T. H., & Otto, M. (2020). The staphylococcal exopolysaccharide PIA – Biosynthesis and role in biofilm formation, colonization, and infection. Computational and Structural Biotechnology Journal, 18, 3324–3334. 10.1016/J.CSBJ.2020.10.027

Nishitani, K., Sutipornpalangkul, W., De Mesy Bentley, K. L., Varrone, J. J., Bello-Irizarry, S. N., Ito, H., Matsuda, S., Kates, S. L., Daiss, J. L., & Schwarz, E. M. (2015). Quantifying the natural history of biofilm formation in vivo during the establishment of chronic implant-associated *Staphylococcus aureus* osteomyelitis in mice to identify critical pathogen and host factors. Journal of orthopaedic research : official publication of the Orthopaedic Research Society, 33(9), 1311–1311. 10.1002/JOR.22907

O’Neill, E., Pozzi, C., Houston, P., Smyth, D., Humphreys, H., Robinson, D. A., & O’Gara, J. P. (2007). Association between methicillin susceptibility and biofilm regulation in *Staphylococcus aureus* isolates from device-related infections. J Clin Microbiol, 45(5), 1379–1388. 10.1128/JCM.02280-06

Olson, M. E., Nygaard, T. K., Ackermann, L., Watkins, R. L., Zurek, O. W., Pallister, K. B., Griffith, S., Kiedrowski, M. R., Flack, C. E., Kavanaugh, J. S., Kreiswirth, B. N., Horswill, A. R., & Voyich, J. M. (2013). *Staphylococcus aureus* nuclease is an SaeRS-dependent virulence factor. Infection and immunity, 81(4), 1316–1324. 10.1128/IAI.01242-12

Paharik, A. E., & Horswill, A. R. (2016). The Staphylococcal Biofilm: Adhesins, Regulation, and Host Response. Microbiology spectrum, 4(2). 10.1128/microbiolspec.VMBF-0022-2015

Peng, D., Chen, A., Shi, B., Min, X., Zhang, T., Dong, Z., Yang, H., Chen, X., Tian, Y., & Chen, Z. (2018). Preliminary study on the effect of brazilin on biofilms of *Staphylococcus aureus*. Experimental and Therapeutic Medicine, 16(3), 2108–2108. 10.3892/ETM.2018.6403

Rice, K. C., Mann, E. E., Endres, J. L., Weiss, E. C., Cassat, J. E., Smeltzer, M. S., & Bayles, K. W. (2007). The *cid*A murein hydrolase regulator contributes to DNA release and biofilm development in *Staphylococcus aureus*. Proceedings of the National Academy of Sciences of the United States of America, 104(19), 8113–8118. 10.1073/PNAS.0610226104

Schurig-Briccio, L. A., Parraga Solorzano, P. K., Lencina, A. M., Radin, J. N., Chen, G. Y., Sauer, J. D., Kehl-Fie, T. E., & Gennis, R. B. (2020). Role of respiratory NADH oxidation in the regulation of *Staphylococcus aureus* virulence. EMBO Rep, 21(5), e45832. 10.15252/embr.201845832

Schwartz, K., Syed, A. K., Stephenson, R. E., Rickard, A. H., & Boles, B. R. (2012). Functional Amyloids Composed of Phenol Soluble Modulins Stabilize *Staphylococcus aureus* Biofilms. PLOS Pathogens, 8(6), e1002744–e1002744. 10.1371/JOURNAL.PPAT.1002744

Seidl, K., Goerke, C., Wolz, C., Mack, D., Berger-Bachi, B., & Bischoff, M. (2008). *Staphylococcus aureus* CcpA affects biofilm formation. Infect Immun, 76(5), 2044–2050. 10.1128/IAI.00035-08

Shopova, I., Bruns, S., Thywissen, A., Kniemeyer, O., Brakhage, A. A., & Hillmann, F. (2013). Extrinsic extracellular DNA leads to biofilm formation and colocalizes with matrix polysaccharides in the human pathogenic fungus *Aspergillus fumigatus*. Frontiers in Microbiology, 4(JUN). 10.3389/FMICB.2013.00141

Sugimoto, S., Sato, F., Miyakawa, R., Chiba, A., Onodera, S., Hori, S., & Mizunoe, Y. (2018). Broad impact of extracellular DNA on biofilm formation by clinically isolated Methicillin-resistant and -sensitive strains of *Staphylococcus aureus*. Scientific reports, 8(1), 2254. 10.1038/s41598-018-20485-z

Tan, L., Huang, Y., Shang, W., Yang, Y., Peng, H., Hu, Z., Wang, Y., Rao, Y., Hu, Q., Rao, X., Hu, X., Li, M., Chen, K., & Li, S. (2022). Accessory Gene Regulator (agr) Allelic Variants in Cognate *Staphylococcus aureus* Strain Display Similar Phenotypes. Frontiers in Microbiology, 13, 700894–700894. 10.3389/FMICB.2022.700894/BIBTEX

Tan, X., Qin, N., Wu, C., Sheng, J., Yang, R., Zheng, B., Ma, Z., Liu, L., Peng, X., & Jia, A. (2015). Transcriptome analysis of the biofilm formed by methicillin-susceptible *Staphylococcus aureus*. Scientific Reports 2015 5:1, 5(1), 1–12. 10.1038/srep11997

Tang, H., Porras, G., Brown, M. M., Chassagne, F., Lyles, J. T., Bacsa, J., Horswill, A. R., & Quave, C. L. (2020). Triterpenoid acids isolated from Schinus terebinthifolia fruits reduce *Staphylococcus aureus* virulence and abate dermonecrosis. Scientific Reports 2020 10:1, 10(1), 1–13. 10.1038/s41598-020-65080-3

Tayeb-Fligelman, E., Tabachnikov, O., Moshe, A., Goldshmidt-Tran, O., Sawaya, M. R., Coquelle, N., Colletier, J. P., & Landau, M. (2017). The cytotoxic *Staphylococcus aureus* PSMα3 reveals a cross-α amyloid-like fibril. Science, 355(6327), 831–833. 10.1126/SCIENCE.AAF4901/SUPPL_FILE/TAYEB-FLIGELMAN.SM.PDF

Trotonda, M. P., Tamber, S., Memmi, G., & Cheung, A. L. (2008). MgrA Represses Biofilm Formation in *Staphylococcus aureus*. Infection and Immunity, 76(12), 5645–5645. 10.1128/IAI.00735-08

Tuchscherr, L., & Otto, M. (2023). Critical Assessment of the Prospects of Quorum-Quenching Therapy for *Staphylococcus aureus* Infection. International Journal of Molecular Sciences 2023, Vol. 24, Page 4025, 24(4), 4025–4025. 10.3390/IJMS24044025

Vij, T., Anil, P. P., Shams, R., Dash, K. K., Kalsi, R., Pandey, V. K., Harsányi, E., Kovács, B., & Shaikh, A. M. (2023). A Comprehensive Review on Bioactive Compounds Found in *Caesalpinia sappan*. Molecules, 28(17). 10.3390/MOLECULES28176247

Vila, T., Kong, E. F., Ibrahim, A., Piepenbrink, K., Shetty, A. C., McCracken, C., Bruno, V., & Jabra-Rizk, M. A. (2019). Candida albicans quorum-sensing molecule farnesol modulates staphyloxanthin production and activates the thiol-based oxidative-stress response in Staphylococcus aureus. Virulence, 10(1), 625–642. 10.1080/21505594.2019.1635418

Vinodhini, V., & Kavitha, M. (2024). Deciphering agr quorum sensing in *Staphylococcus aureu*s: insights and therapeutic prospects. Molecular biology reports, 51(1). 10.1007/S11033-023-08930-3

Wang, H. Y., Hua, X. W., Jia, H. R., Li, C., Lin, F., Chen, Z., & Wu, F. G. (2016). Universal Cell Surface Imaging for Mammalian, Fungal, and Bacterial Cells. ACS Biomater Sci Eng, 2(6), 987–997. 10.1021/acsbiomaterials.6b00130

Xue, L., Chen, Y. Y., Yan, Z., Lu, W., Wan, D., & Zhu, H. (2019). Staphyloxanthin: a potential target for antivirulence therapy. Infection and Drug Resistance, 12, 2151–2151. 10.2147/IDR.S193649

