## Supplementary figures for "Multifaceted suppression of staphylococcal virulence: *in vitro* study of a cellular status with reduced Agr activity and impaired biofilm"

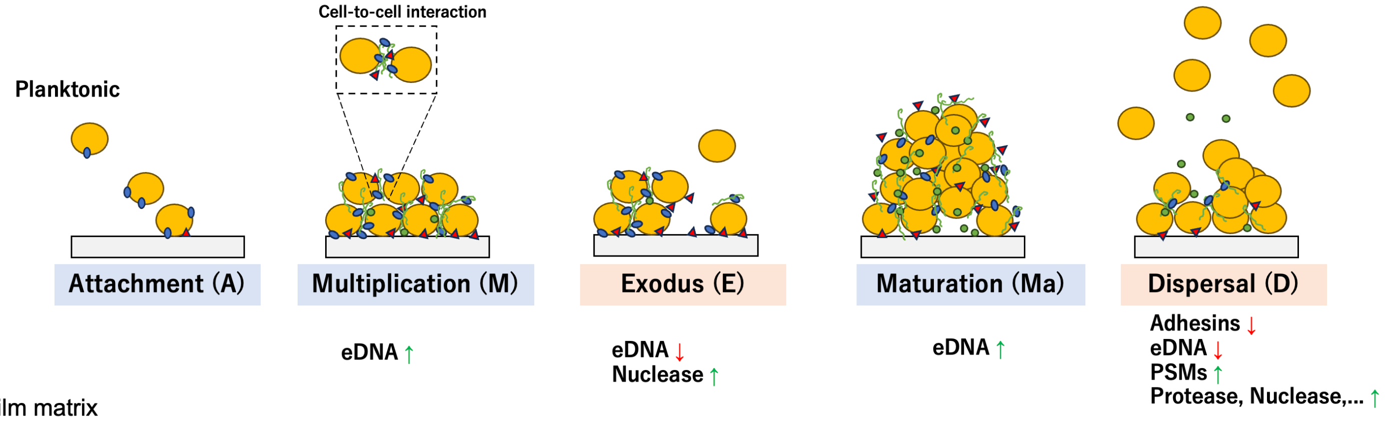


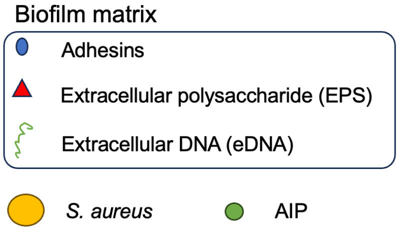


| Stage | A~Ma | A~Ma | E | D | D |
|---|---|---|---|---|---|
| Factors | EPS | Adhersins | Nuclease | Protease | PSMs |
| **SarA** | + | + | | + | |
| **SaeRS** | | + | + | + | |
| **Agr** | | *via* Rot | | + | + |

+ : regulate

***Fig S1.* Biofilm development and factors involved**

Biofilm development and the factors involved are well summarized in review articles (Moormeier & Bayles, 2017; Paharik & Horswill, 2016). In the attachment and multiplication stages, the microbial surface components are involved in cell adherence and cell-to-cell interactions. The involved factors are EPS (PIA) and cell wall-associated proteins such as staphylococcal protein A (SpA), fibronectin-binding proteins (FnBPA, FnBPB), collagen-binding protein, and clumping factors (ClpA, ClpB). The eDNA released by cell death contributes to biofilm stabilization by serving as a "glue" between cells via interaction with extracellular and cell wall-associated proteins and PIA in the biofilm matrix (Campoccia et al., 2021). The exodus stage precedes the maturation of biofilm. The biofilm is reconstructed through the early dispersal by the nuclease (Nuc) expression in the subpopulation that destroys the eDNA. The Nuc expression is under the control of the SaePQRS system (Liu et al., 2016). The exodus stage may contribute to architecting the mature biofilm to enhance nutrient exchange and waste removal (Moormeier et al., 2014). In the dispersal stage, exoenzymes and phenol soluble modulins (PSMs) disseminate cells to distal sites.

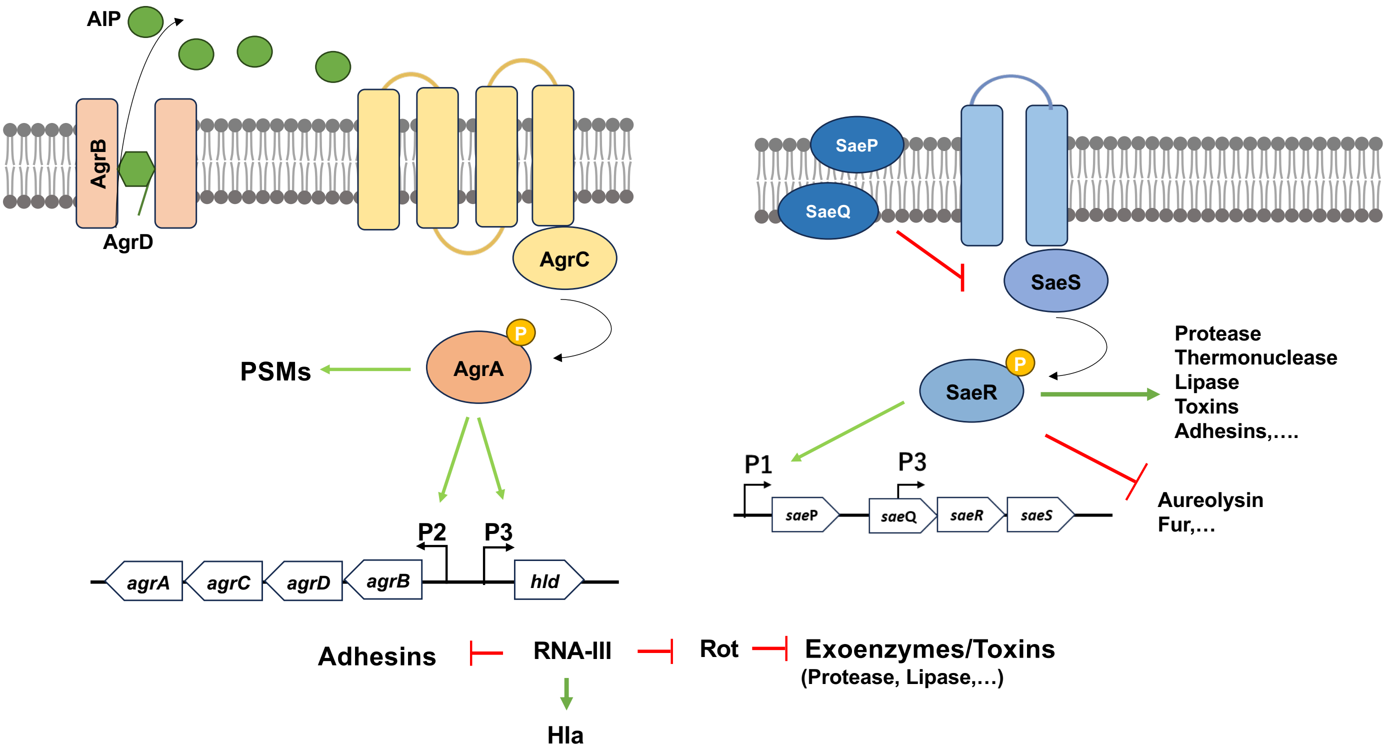


***Fig S2*. Key virulence regulators: Agr system and SaeRS system**

The Agr system is encoded by two regions under the control of P2 and P3 promoters. The *agrBDCA* operon encodes the quorum sensing system. The *agrD* gene encodes the peptide precursor for AIP. AgrD is then modified and exported by the transmembrane endopeptidase AgrB. The *agrC* and *agrA* genes encode the two-component system to sense the AIP concentration that increases in high cell density or closed environment such as in phagosomes (thereby a.k.a diffusion sensing system). The histidine kinase sensor AgrC activates AgrA by phosphorylation. The activated AgrA directly upregulates the P2 and P3 promoters and expression of PSMs. The P3 promoter controls the transcription of the RNA-III factor. RNA-III itself encodes δ-hemolysin (*hld*). By forming RNA duplexes, RNA-III also acts as a translational regulator. It directly or indirectly (*via* Rot) activates the expression of exoenzymes such as nuclease, lipases, and leukocidins, and blocks that of a series of surface proteins (SpA, FnBPs, etc).

The *S. aureus* exoprotein expression (Sae) SaeRS system is the two-component system regulating the expression of virulence factors including toxins and immune-evasive proteins (Liu et al., 2016). SaeRS regulon includes α-, β- and γ-hemolysins, coagulase, Panton-Valentine leucocidin (LukGH), TSST-1, enterotoxin-like toxin, nuclease and adhesins (Eap, FnbAB, Efb, Eab, etc.). The *saePQRS* operon is under the control of two promoters, P1 and P3. Human neutrophil peptides (HNP 1 to 3), calprotectin, hydrogen peroxide, and subinhibitory concentrations of β-lactam antibiotics are reported to activate the SaeRS. On the other hand, several factors, such as low pH conditions, high concentrations of NaCl, or some fatty acids, affect it negatively. The activated SaeR leads to the autoactivation of the promoter P1. SaePQ also controls this autoactivation. Moreover, Agr system and Rot are involved indirectly in the regulation of *saePQRS*.

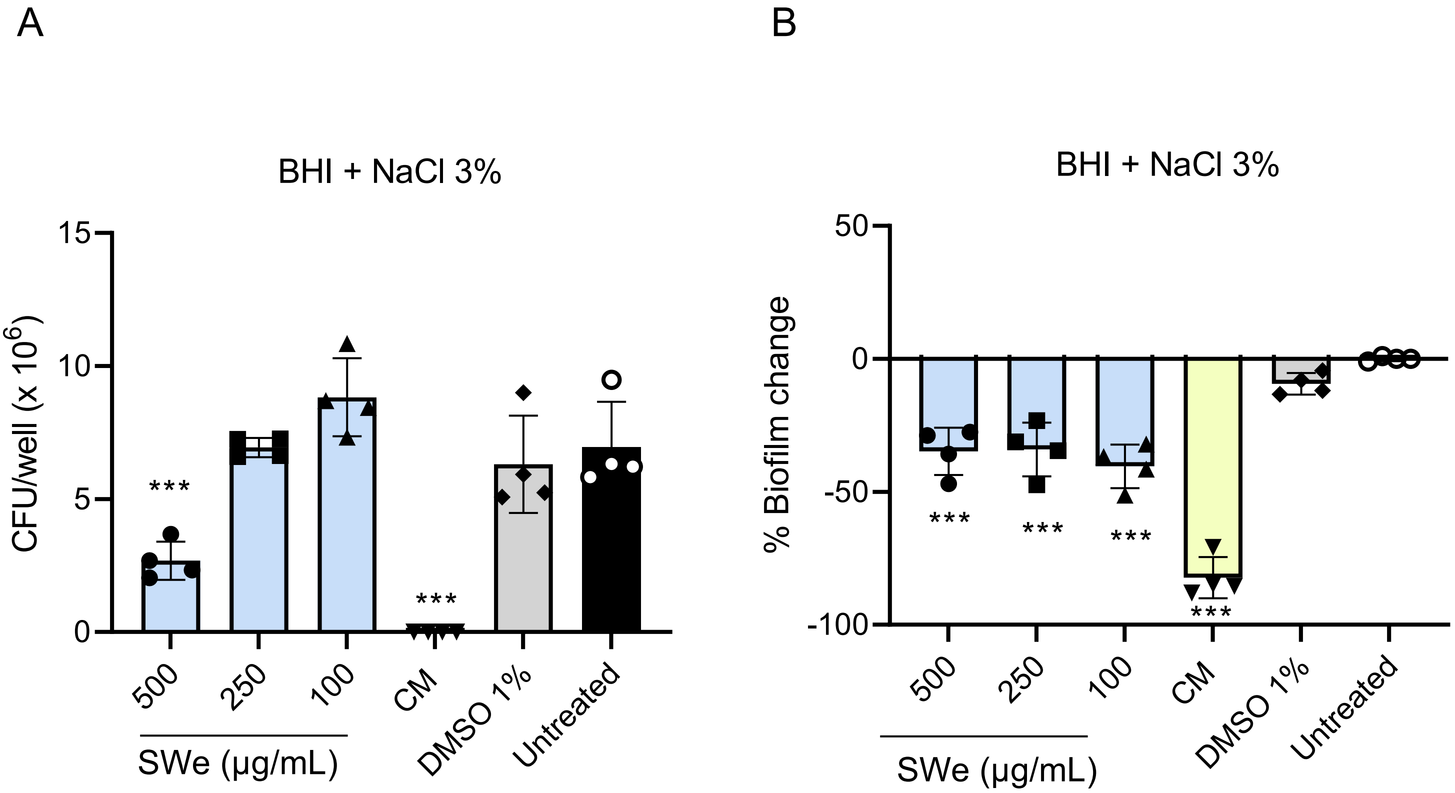


***Fig S3.* The antibiofilm activity of SWe on SH1000 in BHI + NaCl3% (EPS-inducing media) after 24 hours of treatment.** (A) Cell viability in total biomass (biofilm and planktonic cells) relative to the untreated sample. The viable cells were counted as CFU. (B) Biofilm was quantified using crystal violet, and the values of percent change relative to the untreated group are shown. Error bars indicate the SD. *: p≤ 0.05, **: p≤0.001, ***: p≤0.0001 compared to the untreated group. One-way ANOVA & Dunnett's multiple comparison test with the untreated group. CM: 62.5 μg/mL chloramphenicol.

The 250μg/mL SWe reduced biofilm without significant growth inhibition, but 500μg/mL SWe reduced the viability and its antibiofilm activity is not clear.

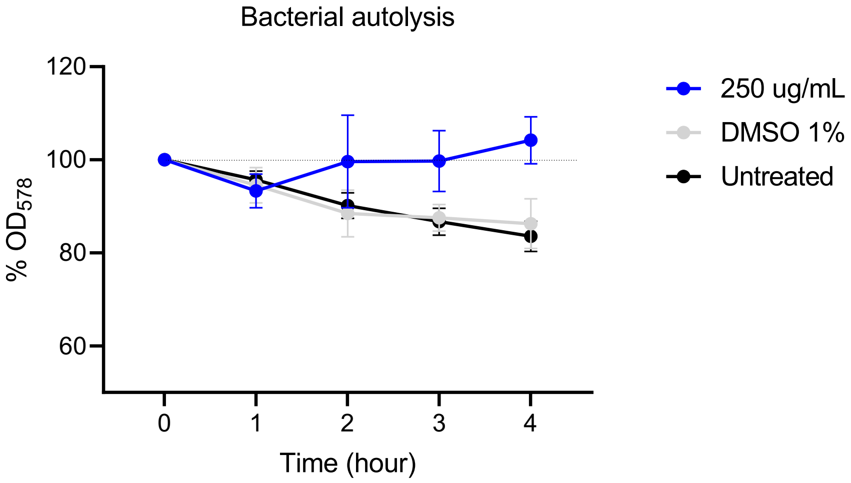


***Fig S4.*** **Autolysis of SH1000 reduced by SWe**.
Cells were pretreated with SWe or 1% DMSO for 3 h, and washed before the assay. The autolysis assay was performed in the shaking condition at 37$^{o}$C in a 96-well plate. $OD_{578}$ was measured by the plate reader.

**Biofilm Disruption assay suggest the qualitative change of the biofilm by SWe$^{lot2}$.**

The first lot of SWe used through this study was not available anymore, and we used a second lot of SWe (SWe$^{lot2}$) for this additional assay. The SWe$^{lot2}$-biofilm was treated by DNase I, Protease K, and metaperiodate that degrade polysaccharides. DNase treatment slightly weakened the pre-formed biofilms irrespective of the presence of SWe$^{lot2}$. On the other hand, inclusion of DNase I throughout the 24h-incubation period reduced the biofilm by about 60% in the absence of SWe$^{lot2}$ (right top well), but it rather increased the biofilm under treatment of 450µg/ml SWe$^{lot2}$. This suggests that in SWe$^{lot2}$ conditions, the role of eDNA for biofilm formation is altered. Protease K reduced the biofilm in normal and SWe$^{lot2}$-biofilm, indicating that protein factors are essential in both cases. Sodium metaperiodate solidified the biofilm (or sedimented cells), and the biofilm in the 450 µg/ml SWe$^{lot2}$ well remained after the washing process; The polysaccharide likely contributes to sustaining the non-biofilm status in the SWe$^{lot2}$-biofilm.

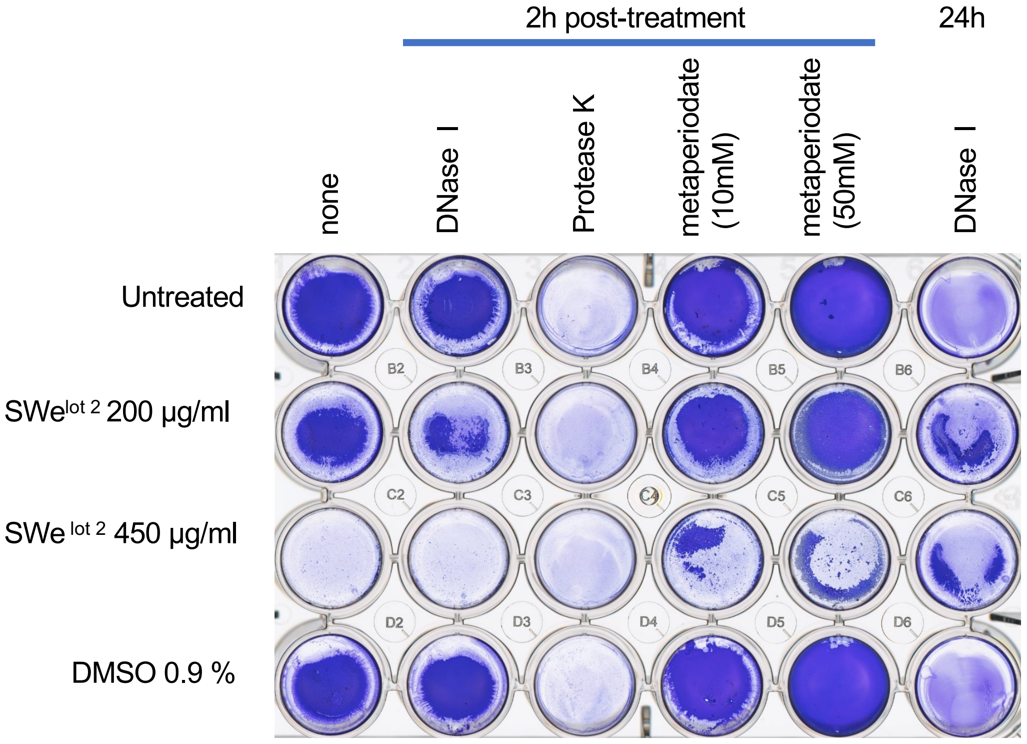


| | | | | | |
|---|---|---|---|---|---|
| **100** | **91**±14 | **28**±8.3 | **104**±7.1 | **133**±4.8 | **39**±14 |
| **84**±4.5 | **69**±6.6 | **32**±2 | **94**±0.6 | **121**±7.2 | **68**±6.7 |
| **32**±4.6 | **28**±9.2 | **37**±1.6 | **62**±9.5 | **98**±27 | **55**±2.9 |
| **103**±18 | **89**±6.4 | **35**±4.5 | **117**±12 | **143**±12 | **36**±14 |

***Fig S5*. Biofilm disruption assay.** Cells were statically cultured with or without SWe$^{lot2}$ (lot number 2) in a 24-well plate. DMSO is the solvent control (0.9%, same with 450µg/ml SWe). After 24h incubation, DNase I, Protease K, or metaperiodate were added directly, without refreshing the medium. In the right column, DNase was added at the beginning and the biofilm was formed in the presence of DNase. Biofilm was quantified by the optimized crystal violet method and normalized to the non-treated biofilm (left top). Mean (bold) and ±SD values are shown (n=2).

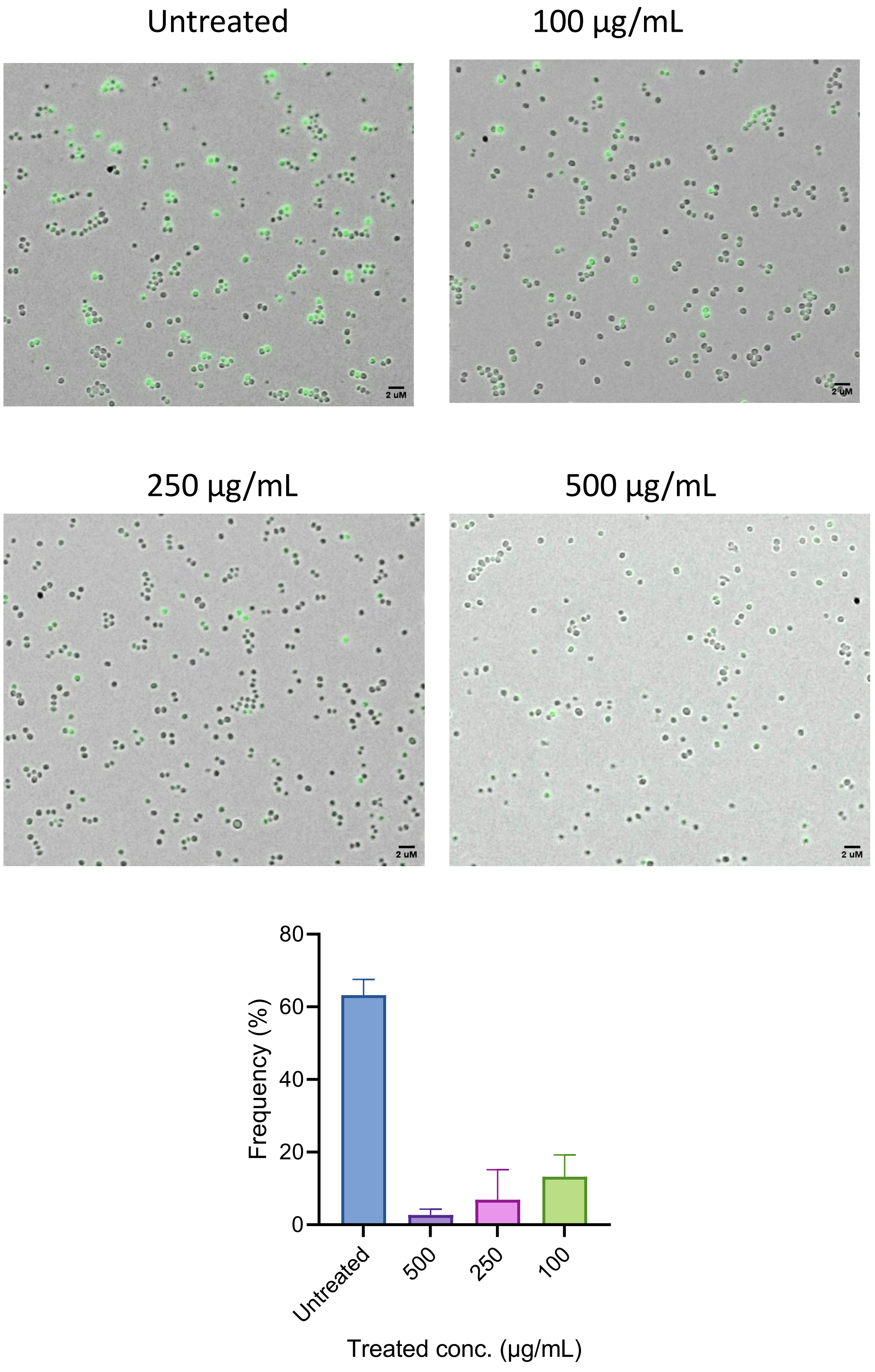


***Fig S6.*** **Microscopy images of SH1000 P3*venus* reporter strain after 20h incubation with SWe.** Fluorescent cells were counted, and the average percentage of fluorescent cells was shown. The error bars show SDs. n=2.

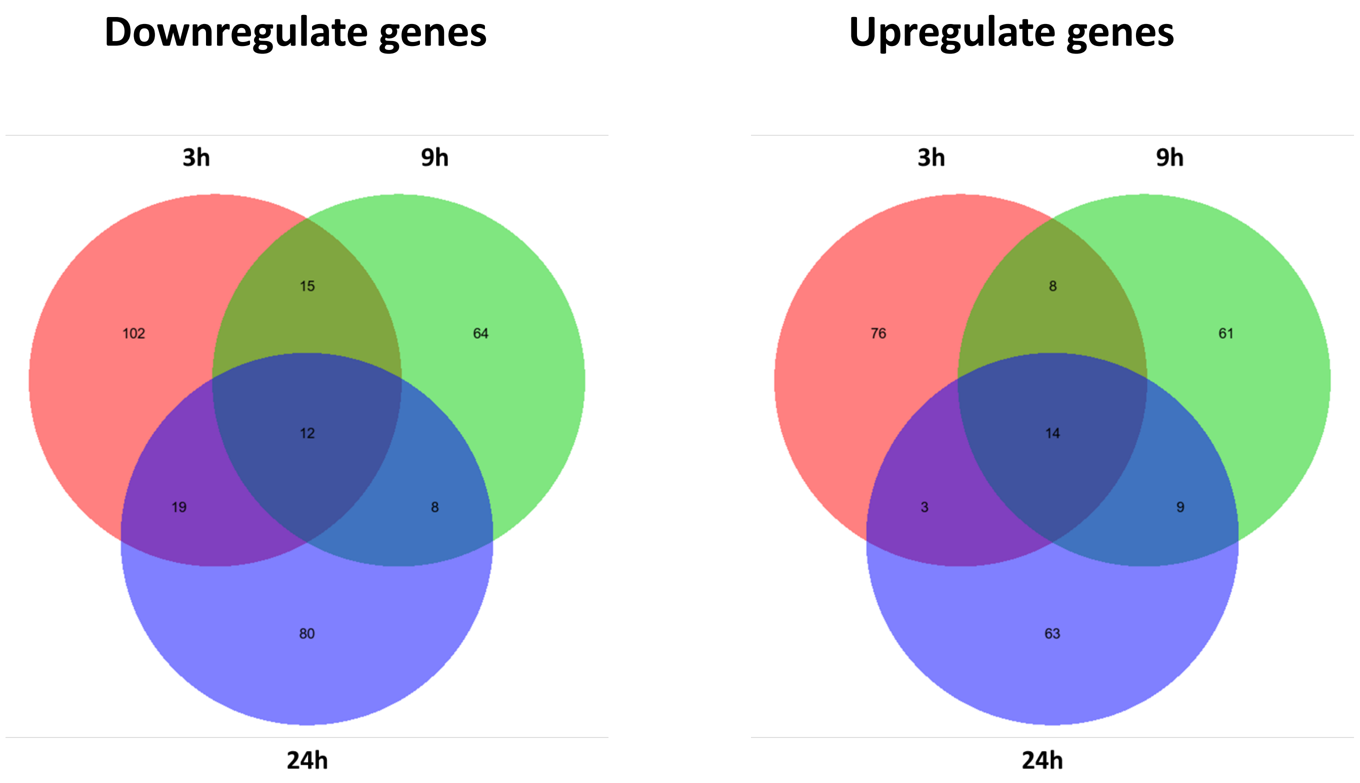


***Fig S7*. Downregulated and upregulated genes shared among 3 h, 9 h, and 24 h.**
Red, green, and blue indicate 3 h, 9 h, and 24 h after the exposure to SWe.

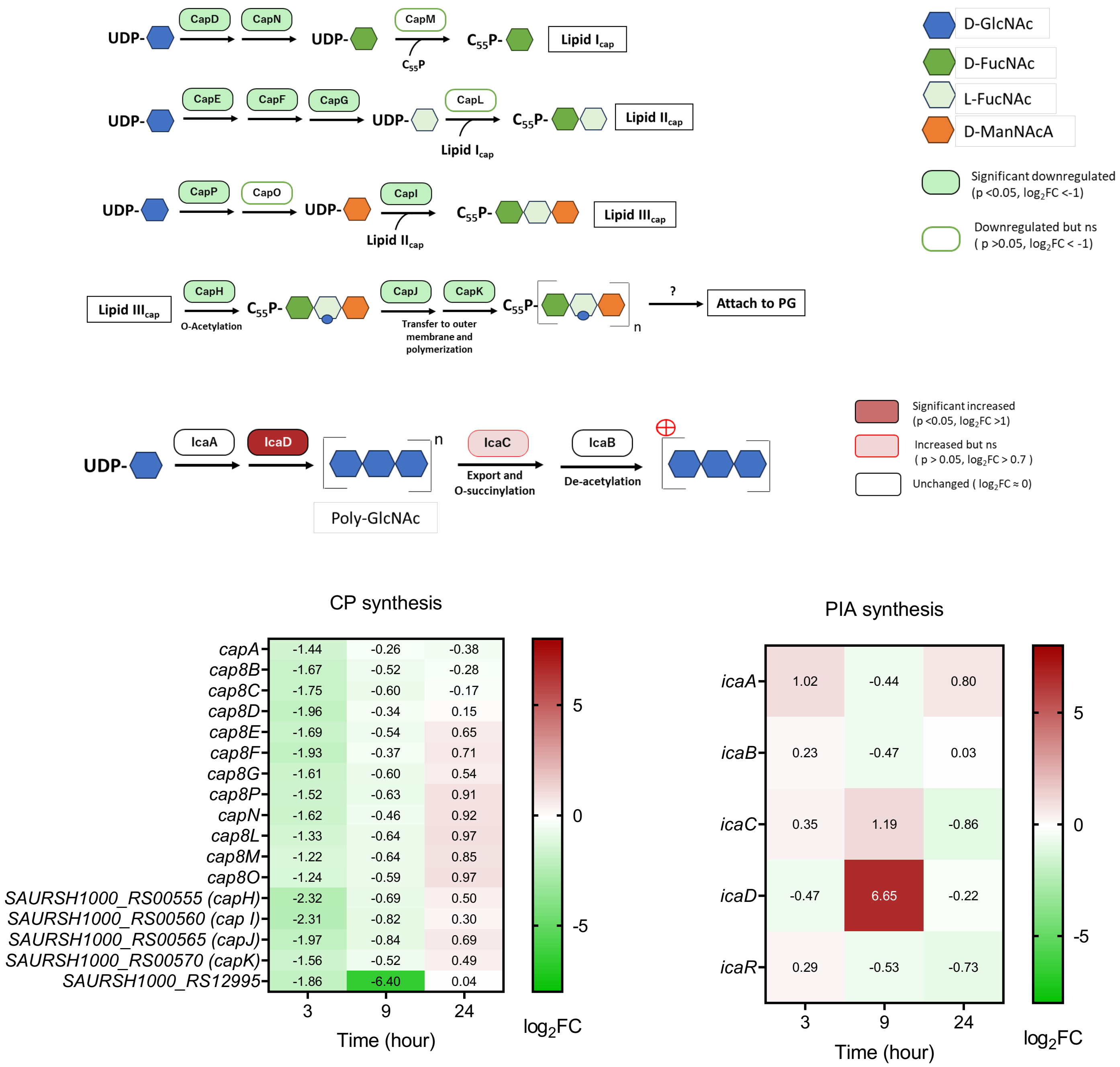


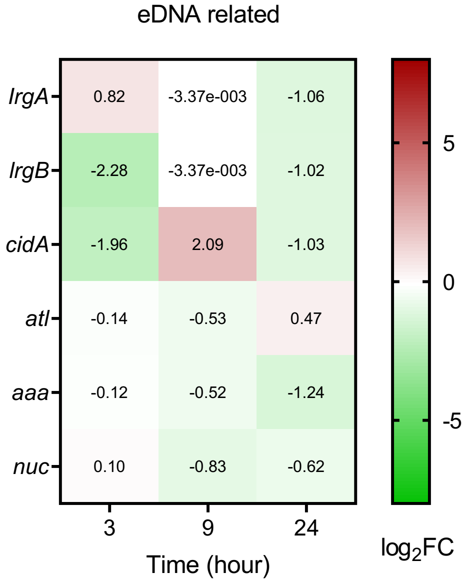


***Fig S8.* DEGs in the SWe-treated cells: polysaccharide- and lysis-related genes.** Synthesis pathways of the capsule and PIA from UDP-GlcNAc are shown top. The UDP-N-acetyl-glucosamine (UDP-N-GlcNAc) is the building material for a series of cell surface components such as PIA, capsule, cell wall, and teichoic acid. Values of $log_2$(Fold Change) relative to untreated control at the same time point are shown. SAURSH1000_RS12995 gene encodes polysaccharide biosynthesis tyrosine autokinase that is involved in the capsule assembly (RefSeq NZ_CP059180.1). C55P: undecaprenyl-phosphate, GlcNAc: N-acetyl-glucosamine, FucNAc: N-acetyl-fucosamine, ManNAcA: N-acetyl-mannosaminuronic acid.

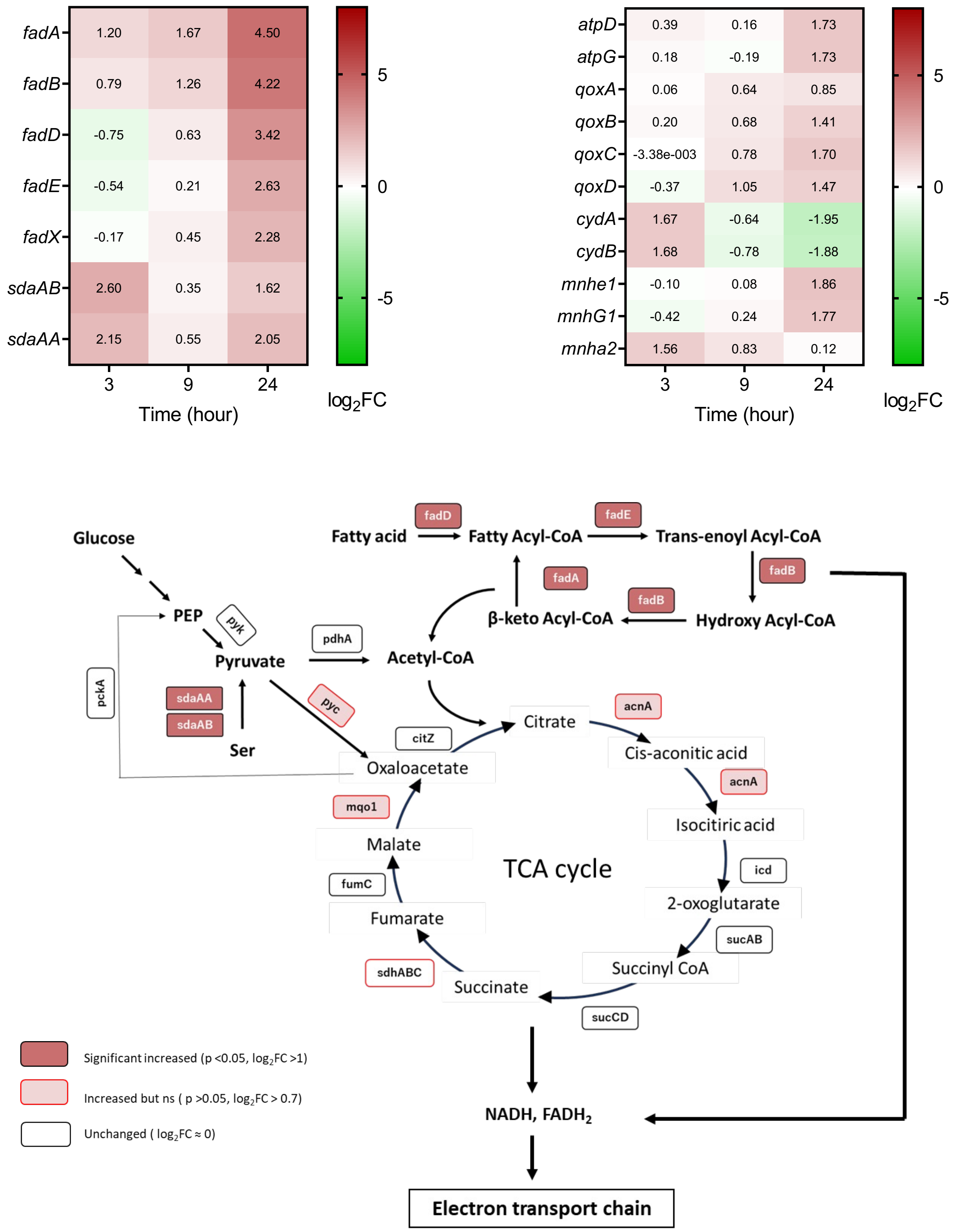


***Fig S9.* Metabolic gene expression affected by SWe.** DEGs related to the catabolism and respiration are shown. The genes coding for fatty acid β-oxidation enzymes (*fadXDEBA*) were increased by SWe at 9 h and 24 h. These enzymes are responsible for the supplementation of acetyl-CoA, which enters the TCA cycle. The gene coding for SdaAA and SdaAB enzyme converting serine to pyruvate were upregulated in 3 h and 24 h. In terms of electron transport, the expression of cytochrome bd oxidase (*cydAB*) was transiently increased at 3 h, and the terminal oxidases *qoxCD*, the Na+/H+ antiporter (*mnhE1*, *mnhG1*) and ATP synthase (*atpCD*) increased at 24 h. These results suggest that cells treated with SWe would have an altered respiration status, which might lead to a change in reactive oxygen species (ROS) level. Indeed, we observed the increase of ROS by SWe at 9 h (**Fig S10**).

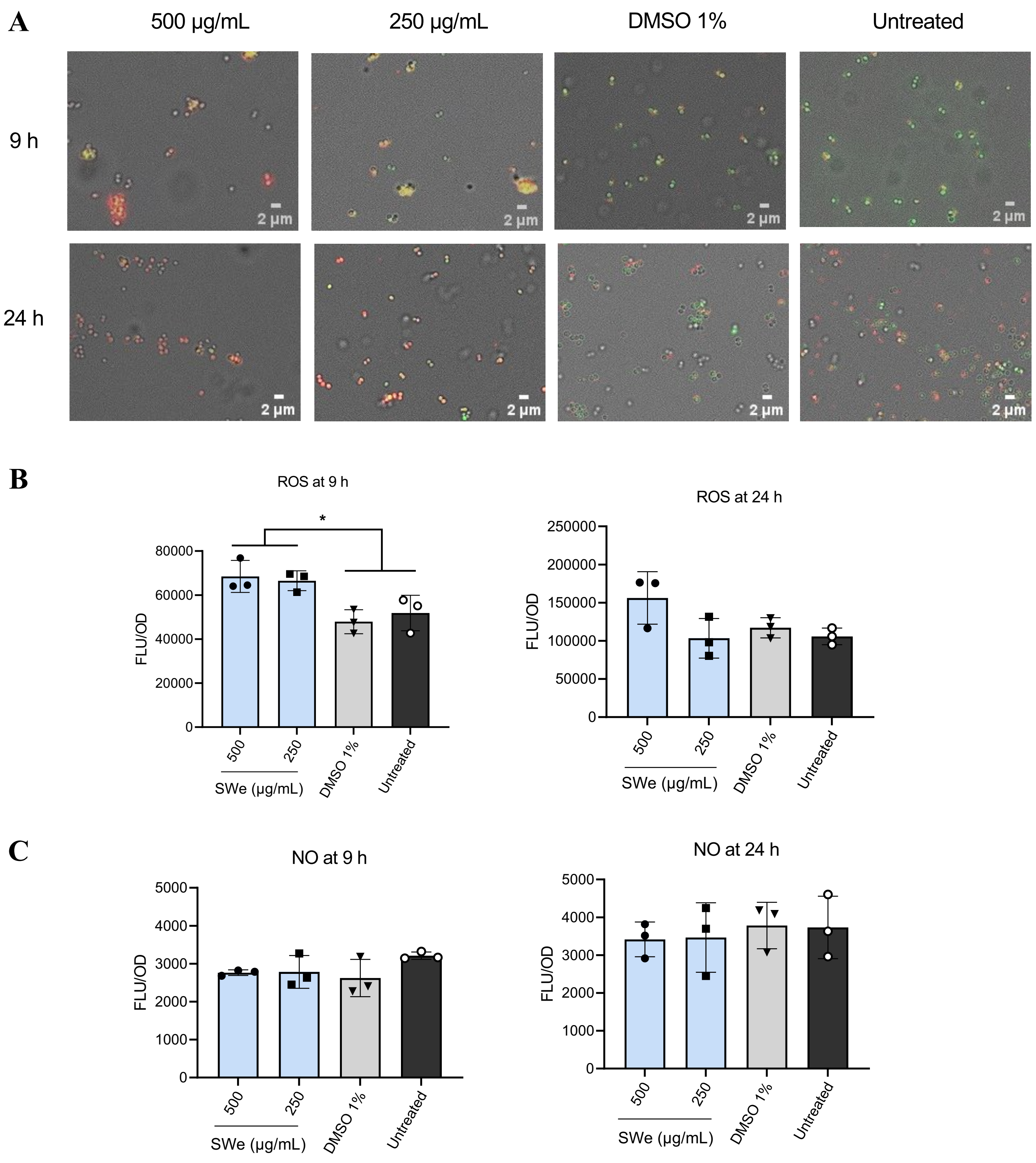


***Fig S10.*** **Intracellular ROS, but not NO, is increased by SWe**. After growing biofilm at indicated times, cells in planktonic and biofilm were collected together as a mixture and stained with CellROX™ (red) and DAF-FM (Green) for detecting the intracellular ROS and NO, respectively. (A) Representative microscopic images. (B) Quantities of ROS measured by CellROX™ and (C) NO detected by DAF-FM at 9 h and 24 h of total biomass. All data represent the average from 3 independent experiments; error bars indicate the SD. * *p≤ 0.05*. One-way ANOVA & Dunnett's multiple comparison test with the untreated group.

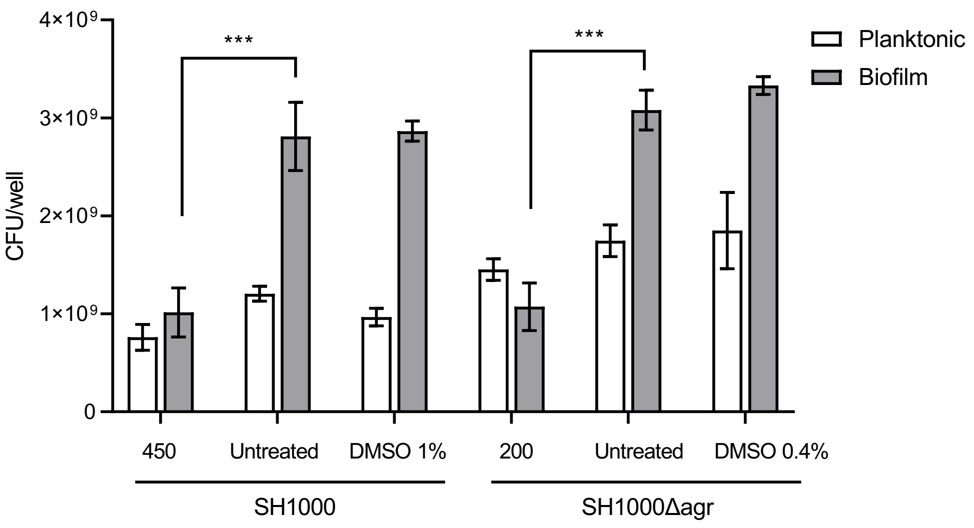


**Fig S11. Anti-biofilm activity of SWe$^{lot2}$ does not depend on *agr*.**
SWe$^{lot2}$ was used instead of SWe that was not available anymore. The appropriate concentration to observe the antibiofilm activity of SWe$^{lot2}$ was 450µg/mL for SH1000 and 200µg/mL for SH1000Δagr. Higher concentration of SWe$^{lot2}$ (e.g. 450µg/mL) had a growth-inhibiting effect on SH1000Δagr and not shown here. Concentrations of DMSO solvent controls are equivalent to the tested SWe$^{lot2}$. Biofilm was formed in a 24-well plate. SWe$^{lot2}$ reduced the biofilm CFU (filled bars), while hardly affecting the planktonic CFU (blank bars), suggesting that anti-biofilm activity does not rely on the presence of the Agr system. Two-way ANOVA & Dunnett's multiple comparison test with the untreated group for each strain (***: *p≤0.0001*).

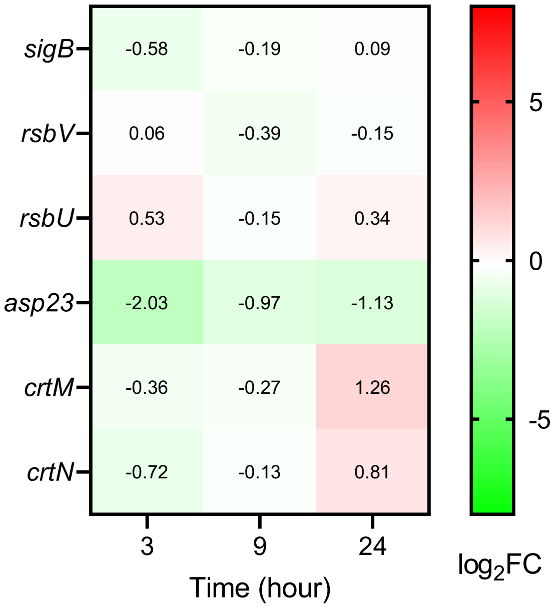


***Fig S12.*** **DEGs in the SWe-treated cells: SigB related genes.**

SigB activity is post-translationally regulated by anti-sigma factor RsbW, and upstream regulators RsbV and RsbU. The mRNA levels of these genes were not affected by SWe. However, the *asp23* gene expression was reduced by SWe, suggesting that SigB activity is reduced by SWe. *crtMN* genes are responsible for carotenoid synthesis (see Discussion).
