## Supplementary tables for "Multifaceted suppression of staphylococcal virulence: *in vitro* study of a cellular status with reduced Agr activity and impaired biofilm"

**Table S1.** MIC values (μg/mL) of SWe in MHB and TSB

| | **RN4220** | **MW2** | **SH1000** | **ATCC25923** |
|---|---|---|---|---|
| **MHB** | 125 | 125 | 500 | 125 |
| **TSB** | 250 | 250 | 500 | 125 |

**Table S2.** DEGs of SWe-treated *S. aureus* compared to negative control at 3 h ($p$ <0.05)

| Gene | Description | Log2FC |
|---|---|---|
| | | 3h |
| cstR | persulfide-sensing transcriptional repressor CstR | 5.10 |
| carA | carbamoyl phosphate synthase small subunit | 4.37 |
| carB | carbamoyl-phosphate synthase large subunit | 4.33 |
| pyrE | orotate phosphoribosyltransferase | 5.40 |
| pyrF | orotidine-5'-phosphate decarboxylase | 4.31 |
| pyrR | bifunctional pyr operon transcriptional regulator/uracil phosphoribosyltransferase PyrR | 1.64 |
| hisD | histidinol dehydrogenase | 3.25 |
| hisH | imidazole glycerol phosphate synthase subunit HisH | 2.34 |
| hisB | imidazoleglycerol-phosphate dehydratase HisB | 5.10 |
| hisG | ATP phosphoribosyltransferase | 4.98 |
| argG | argininosuccinate synthase | 3.13 |
| argH | argininosuccinate lyase | 2.55 |
| sdaaB | L-serine ammonia-lyase, iron-sulfur-dependent subunit beta | 2.60 |
| leuD | 3-isopropylmalate dehydratase small subunit | 2.19 |
| glpk | glycerol kinase GlpK | 2.14 |
| trep | PTS system trehalose-specific EIIBC component | 2.14 |
| dapB | 4-hydroxy-tetrahydrodipicolinate reductase | 2.09 |
| dapD | 2,3,4,5-tetrahydropyridine-2,6-dicarboxylate N-acetyltransferase | 1.81 |
| nrdD | anaerobic ribonucleoside-triphosphate reductase | 1.71 |
| mepa | multidrug efflux MATE transporter MepA | 1.66 |
| dapA | 4-hydroxy-tetrahydrodipicolinate synthase | 2.37 |
| quec | 7-cyano-7-deazaguanine synthase QueC | 2.34 |
| ilvc | ketol-acid reductoisomerase | 1.62 |
| tgt | tRNA guanosine(34) transglycosylase Tgt | 1.58 |
| mnha2 | Na+/H+ antiporter Mnh2 subunit A | 1.56 |
| arsb | arsenite efflux transporter membrane subunit ArsB | 1.55 |
| vraH | peptide resistance ABC transporter activity modulator VraH | 4.69 |
| comGB | competence type IV pilus assembly protein ComGB | 5.10 |
| SAURSH1000_RS01850 | superantigen-like protein SSL9 | 6.39 |
| SAURSH1000_RS00220 | LysR family transcriptional regulator | 5.88 |
| SAURSH1000_RS11090 | hypothetical protein | 5.67 |
| SAURSH1000_RS12720 | hypothetical protein | 5.41 |
| SAURSH1000_RS06250 | hypothetical protein | 5.10 |
| SAURSH1000_RS02115 | 23S ribosomal RNA | 4.69 |
| SAURSH1000_RS02360 | tRNA-Ala | 4.69 |
| SAURSH1000_RS04600 | YxeA family protein | 4.69 |
| SAURSH1000_RS09725 | ACT domain-containing protein | 4.18 |
| SAURSH1000_RS13035 | hypothetical protein | 4.03 |
| SAURSH1000_RS05465 | solute carrier family 23 protein | 4.02 |
| SAURSH1000_RS05475 | dihydroorotase | 3.99 |
| SAURSH1000_RS03550 | ABC transporter ATP-binding protein | 3.85 |
| SAURSH1000_RS05470 | aspartate carbamoyltransferase catalytic subunit | 3.66 |
| SAURSH1000_RS01315 | TIGR01741 family protein | 3.64 |
| SAURSH1000_RS04145 | Na+/H+ antiporter family protein | 3.16 |
| SAURSH1000_RS06495 | aspartate kinase | 3.06 |
| SAURSH1000_RS01370 | winged helix-turn-helix transcriptional regulator | 2.97 |
| SAURSH1000_RS06500 | aspartate-semialdehyde dehydrogenase | 2.57 |
| SAURSH1000_RS13075 | ATP phosphoribosyltransferase regulatory subunit | 2.49 |
| SAURSH1000_RS01450 | NAD-dependent deacetylase | 2.34 |

| | | |
|---|---|---|
| SAURSH1000_RS00815 | maltodextrin ABC transporter substrate-binding protein | 2.28 |
| SAURSH1000_RS11800 | NAD(P)-dependent oxidoreductase | 2.26 |
| SAURSH1000_RS09600 | nitroreductase family protein | 2.17 |
| SAURSH1000_RS00270 | M20 family metallopeptidase | 2.06 |
| SAURSH1000_RS12485 | PTS transporter subunit IIC | 2.00 |
| SAURSH1000_RS03545 | iron chelate uptake ABC transporter family permease subunit | 1.96 |
| SAURSH1000_RS11170 | hypothetical protein | 1.95 |
| SAURSH1000_RS08925 | ABC transporter permease subunit | 1.92 |
| SAURSH1000_RS01525 | GNAT family protein | 1.90 |
| SAURSH1000_RS10260 | PTS mannitol transporter subunit IICB | 1.90 |
| SAURSH1000_RS12410 | sugar O-acetyltransferase | 1.89 |
| SAURSH1000_RS01395 | ROK family protein | 1.84 |
| SAURSH1000_RS04610 | hypothetical protein | 1.83 |
| SAURSH1000_RS03240 | GNAT family N-acetyltransferase | 1.78 |
| SAURSH1000_RS00820 | sugar ABC transporter permease | 1.75 |
| SAURSH1000_RS02860 | aldo/keto reductase | 1.75 |
| SAURSH1000_RS10405 | YjiH family protein | 1.69 |
| SAURSH1000_RS04885 | cytochrome d ubiquinol oxidase subunit II | 1.68 |
| SAURSH1000_RS04880 | cytochrome ubiquinol oxidase subunit I | 1.67 |
| SAURSH1000_RS09230 | hypothetical protein | 1.66 |
| SAURSH1000_RS13110 | YceI family protein | 1.61 |
| SAURSH1000_RS00085 | adenylosuccinate synthase | 1.60 |
| SAURSH1000_RS03540 | ABC transporter permease | 1.59 |
| SAURSH1000_RS06940 | YpdA family putative bacillithiol disulfide reductase | 1.56 |
| SAURSH1000_RS07805 | hypothetical protein | 1.56 |
| SAURSH1000_RS11045 | CHAP domain-containing protein | 1.50 |
| SAURSH1000_RS01335 | formate/nitrite transporter family protein | 1.45 |
| isdD | iron-regulated surface determinant protein IsdD | -6.29 |
| fdh | NAD-dependent formate dehydrogenase | -3.95 |
| arcA | arginine deiminase | -2.69 |
| arcB | ornithine carbamoyltransferase | -3.62 |
| arcC | carbamate kinase | -2.18 |
| arcD | arginine-ornithine antiporter | -2.20 |
| agrA | LytTR family DNA-binding domain-containing protein | -2.77 |
| agrB | accessory gene regulator AgrB | -2.71 |
| agrC | GHKL domain-containing protein | -3.06 |
| agrD | cyclic lactone autoinducer peptide | -3.71 |
| phnC | phosphonate ABC transporter ATP-binding protein | -2.41 |
| essB | type VII secretion protein EssB | -2.32 |
| cidA | holin-like murein hydrolase modulator CidA | -1.96 |
| lrgB | antiholin-like protein LrgB | -2.28 |
| graF | glycopeptide resistance-associated protein GraF | -2.24 |
| fnbB | fibronectin-binding protein FnbB | -2.16 |
| ald | alanine dehydrogenase | -2.14 |
| clpL | ATP-dependent Clp protease ATP-binding subunit ClpL | -2.02 |
| isdA | LPXTG-anchored heme-scavenging protein IsdA | -1.88 |
| ebpS | elastin-binding protein EbpS | -1.80 |
| norB | multidrug efflux MFS transporter NorB | -1.79 |
| lip2 | YSIRK domain-containing triacylglycerol lipase Lip2/Geh | -1.77 |
| tdcB | bifunctional threonine ammonia-lyase/L-serine ammonia-lyase TdcB | -1.75 |
| fmhA | FemA/FemB family glycyltransferase FmhA | -1.57 |
| vwB | von Willebrand factor binding protein Vwb | -2.73 |
| sarS | HTH-type transcriptional regulator SarS | -1.65 |

| | | |
|---|---|---|
| capA | capsular polysaccharide type 5/8 biosynthesis protein CapA | -1.44 |
| cap8B | type 8 capsular polysaccharide synthesis protein Cap8B | -1.67 |
| cap8C | type 8 capsular polysaccharide synthesis protein Cap8C | -1.75 |
| cap8D | type 8 capsular polysaccharide synthesis protein Cap8D | -1.96 |
| cap8E | type 8 capsular polysaccharide synthesis protein Cap8E | -1.69 |
| cap8F | type 8 capsular polysaccharide synthesis protein Cap8F | -1.93 |
| cap8G | type 8 capsular polysaccharide synthesis protein Cap8G | -1.61 |
| cap8P | type 8 capsular polysaccharide synthesis protein Cap8P | -1.52 |
| capN | capsular polysaccharide type 5/8 biosynthesis epimerase CapN | -1.62 |
| SAURSH1000_RS00570 | capsular polysaccharide biosynthesis protein | -1.56 |
| SAURSH1000_RS06060 | minor capsid protein | -1.94 |
| SAURSH1000_RS00555 | CatB-related O-acetyltransferase | -2.32 |
| SAURSH1000_RS00565 | O-antigen ligase | -1.97 |
| SAURSH1000_RS01660 | tyrosine-type recombinase/integrase | -6.29 |
| SAURSH1000_RS07300 | YqgQ family protein | -5.97 |
| SAURSH1000_RS01285 | DUF5080 family protein | -5.56 |
| SAURSH1000_RS10600 | hypothetical protein | -5.56 |
| SAURSH1000_RS01295 | TIGR01741 family protein | -5.31 |
| SAURSH1000_RS04080 | hypothetical protein | -5.31 |
| SAURSH1000_RS06990 | DUF1672 domain-containing protein | -5.31 |
| SAURSH1000_RS02365 | 16S ribosomal RNA | -5.07 |
| SAURSH1000_RS06075 | hypothetical protein | -5.00 |
| SAURSH1000_RS08855 | hypothetical protein | -5.00 |
| SAURSH1000_RS09770 | 16S ribosomal RNA | -4.11 |
| SAURSH1000_RS06980 | DUF1672 domain-containing protein | -3.34 |
| SAURSH1000_RS09090 | 16S ribosomal RNA | -3.28 |
| SAURSH1000_RS11050 | hypothetical protein | -2.89 |
| SAURSH1000_RS03020 | metal ABC transporter ATP-binding protein | -2.79 |
| SAURSH1000_RS01345 | 5'-nucleotidase, lipoprotein e(P4) family | -2.75 |
| SAURSH1000_RS06045 | hypothetical protein | -2.72 |
| SAURSH1000_RS06135 | hypothetical protein | -2.67 |
| SAURSH1000_RS00425 | superoxide dismutase | -2.65 |
| SAURSH1000_RS03010 | zinc ABC transporter substrate-binding protein | -2.64 |
| SAURSH1000_RS00485 | hypothetical protein | -2.45 |
| SAURSH1000_RS07815 | hypothetical protein | -2.32 |
| SAURSH1000_RS00560 | glycosyltransferase | -2.31 |
| SAURSH1000_RS05915 | hypothetical protein | -2.31 |
| SAURSH1000_RS01915 | hypothetical protein | -2.31 |
| SAURSH1000_RS00140 | hypothetical protein | -2.31 |
| SAURSH1000_RS03805 | hypothetical protein | -2.26 |
| SAURSH1000_RS03735 | DUF4887 domain-containing protein | -2.20 |
| SAURSH1000_RS09050 | tRNA-Leu | -2.19 |
| SAURSH1000_RS11730 | ABC transporter ATP-binding protein | -2.19 |
| SAURSH1000_RS00135 | hypothetical protein | -2.09 |
| SAURSH1000_RS05820 | YlxQ family RNA-binding protein | -2.07 |
| SAURSH1000_RS06085 | hypothetical protein | -2.03 |
| SAURSH1000_RS10455 (asp23) | Asp23/Gls24 family envelope stress response protein | -2.03 |
| SAURSH1000_RS03015 | metal ABC transporter permease | -2.02 |
| SAURSH1000_RS08470 | hypothetical protein | -2.02 |
| SAURSH1000_RS04580 | hypothetical protein | -2.01 |
| SAURSH1000_RS01410 | YjiH family protein | -2.00 |
| SAURSH1000_RS00235 | hypothetical protein | -1.98 |
| SAURSH1000_RS10560 | MerR family transcriptional regulator | -1.96 |

| | | |
|---|---|---|
| SAURSH1000_RS11480 | DUF4889 domain-containing protein | -1.91 |
| SAURSH1000_RS02190 | Veg family protein | -1.90 |
| SAURSH1000_RS05535 | hypothetical protein | -1.84 |
| SAURSH1000_RS10615 | AAA family ATPase | -1.79 |
| SAURSH1000_RS05945 | RicAFT regulatory complex protein RicA family protein | -1.77 |
| SAURSH1000_RS00650 | DUF2294 domain-containing protein | -1.75 |
| SAURSH1000_RS07210 | hypothetical protein | -1.70 |
| SAURSH1000_RS11890 | DUF1307 domain-containing protein | -1.67 |
| SAURSH1000_RS10220 | hypothetical protein | -1.62 |
| SAURSH1000_RS02390 | 23S ribosomal RNA | -1.61 |
| SAURSH1000_RS13170 | HdeD family acid-resistance protein | -1.61 |
| SAURSH1000_RS06720 | amino acid permease | -1.61 |
| SAURSH1000_RS09000 | tRNA-Ser | -1.59 |
| SAURSH1000_RS11410 | hypothetical protein | -1.57 |
| SAURSH1000_RS10215 | hypothetical protein | -1.56 |
| SAURSH1000_RS10470 | BCCT family transporter | -1.55 |
| SAURSH1000_RS11330 | TetR/AcrR family transcriptional regulator | -1.53 |
| SAURSH1000_RS12745 | hypothetical protein | -1.53 |
| SAURSH1000_RS12670 | hypothetical protein | -1.50 |
| SAURSH1000_RS11365 | ABC transporter ATP-binding protein | -1.49 |
| SAURSH1000_RS10460 | DUF2273 domain-containing protein | -1.44 |

**Table S3.** DEGs of SWe-treated S. aureus compared to negative control at 9 h (*p* <0.05)

| Gene | Description | Log2FC |
|---|---|---|
| | | 9h |
| icaD | intracellular adhesion protein IcaD | 6.65 |
| purE | 5-(carboxyamino)imidazole ribonucleotide mutase | 2.89 |
| isdF | hemin ABC transporter permease protein IsdF | 2.49 |
| dprA | DNA-processing protein DprA | 2.25 |
| cidA | holin-like murein hydrolase modulator CidA | 2.09 |
| lukE | bi-component leukocidin LukED subunit E | 1.93 |
| pfkB | 1-phosphofructokinase | 1.83 |
| ffs | signal recognition particle sRNA large type | 1.83 |
| esxA | WXG100 family type VII secretion effector EsxA | 1.82 |
| rbsD | D-ribose pyranase | 1.44 |
| czrB | CDF family zinc efflux transporter CzrB | 1.38 |
| folP | dihydropteroate synthase | 1.31 |
| sarX | HTH-type transcriptional regulator SarX | 1.30 |
| pepa1 | type I toxin-antitoxin system Fst family toxin PepA1 | 1.15 |
| qoxD | cytochrome aa3 quinol oxidase subunit IV | 1.05 |
| cwrA | cell wall inhibition responsive protein CwrA | 1.02 |
| SAURSH1000_RS09765 | 23S ribosomal RNA | 6.54 |
| SAURSH1000_RS02840 | DUF443 family protein | 6.13 |
| SAURSH1000_RS08775 | tRNA-Met | 5.56 |
| SAURSH1000_RS09500 | hypothetical protein | 5.56 |
| SAURSH1000_RS12385 | ferrous iron transport protein A | 5.56 |
| SAURSH1000_RS08600 | hypothetical protein | 5.30 |
| SAURSH1000_RS00715 | hypothetical protein | 4.99 |
| SAURSH1000_RS08705 | gallidermin/nisin family lantibiotic | 4.99 |
| SAURSH1000_RS11585 | hypothetical protein | 4.99 |
| SAURSH1000_RS11365 | ABC transporter ATP-binding protein | 3.54 |
| SAURSH1000_RS07815 | hypothetical protein | 3.38 |
| SAURSH1000_RS11370 | ABC transporter permease | 3.06 |
| SAURSH1000_RS04305 | LysR family transcriptional regulator | 2.90 |
| SAURSH1000_RS04570 | YkvS family protein | 2.90 |
| SAURSH1000_RS08540 | DUF4909 domain-containing protein | 2.90 |
| SAURSH1000_RS03350 | DeoR/GlpR family DNA-binding transcription regulator | 2.88 |
| SAURSH1000_RS11995 | hypothetical protein | 2.74 |
| SAURSH1000_RS13060 | histidinol-phosphate transaminase | 2.74 |
| SAURSH1000_RS09000 | tRNA-Ser | 2.65 |
| SAURSH1000_RS09655 | ammonium transporter | 2.51 |
| SAURSH1000_RS13265 | hypothetical protein | 2.39 |
| SAURSH1000_RS01735 | hypothetical protein | 2.38 |
| SAURSH1000_RS01440 | glycine cleavage system protein H | 2.15 |
| SAURSH1000_RS09050 | tRNA-Leu | 2.13 |
| SAURSH1000_RS03360 | fructose-specific PTS transporter subunit EIIC | 1.98 |
| SAURSH1000_RS08940 | tRNA-Leu | 1.95 |
| SAURSH1000_RS10215 | hypothetical protein | 1.88 |
| SAURSH1000_RS01890 | tandem-type lipoprotein | 1.85 |
| SAURSH1000_RS12555 | NAD(P)-binding domain-containing protein | 1.79 |
| SAURSH1000_RS06040 | hypothetical protein | 1.75 |
| SAURSH1000_RS01175 | DUF5080 family protein | 1.70 |

| | | |
|---|---|---|
| SAURSH1000_RS11550 | nitrate reductase subunit alpha | 1.61 |
| SAURSH1000_RS06135 | hypothetical protein | 1.60 |
| SAURSH1000_RS01160 | ABC transporter permease | 1.50 |
| SAURSH1000_RS09365 | NETI motif-containing protein | 1.45 |
| SAURSH1000_RS10415 | iron ABC transporter permease | 1.43 |
| SAURSH1000_RS01385 | sodium:solute symporter | 1.41 |
| SAURSH1000_RS00960 | PTS transporter subunit EIIC | 1.29 |
| SAURSH1000_RS12530 | SDR family NAD(P)-dependent oxidoreductase | 1.25 |
| SAURSH1000_RS02720 | DUF423 domain-containing protein | 1.23 |
| SAURSH1000_RS11445 | YhgE/Pip domain-containing protein | 1.17 |
| SAURSH1000_RS01760 | nucleobase:cation symporter-2 family protein | 1.15 |
| SAURSH1000_RS10220 | hypothetical protein | 1.13 |
| SAURSH1000_RS11505 | nitrate/nitrite transporter | 1.11 |
| SAURSH1000_RS02675 | MFS transporter | 1.06 |
| sarT | HTH-type transcriptional regulator SarT | -5.11 |
| sarU | HTH-type transcriptional regulator SarU | -5.11 |
| sel26 | staphylococcal enterotoxin type 26 | -4.70 |
| comGB | competence type IV pilus assembly protein ComGB | -3.29 |
| splE | serine protease SplE | -2.00 |
| sph (toxin) | sphingomyelin phosphodiesterase | -1.81 |
| lpl9 | tandem-type lipoprotein Lpl9 | -1.76 |
| tilS | tRNA lysidine(34) synthetase TilS | -1.76 |
| hemW | radical SAM family heme chaperone HemW | -1.74 |
| isdB | heme uptake protein IsdB | -1.70 |
| phnE | phosphonate ABC transporter, permease protein PhnE | -1.63 |
| sigS | RNA polymerase sigma factor SigS | -1.44 |
| recX | recombination regulator RecX | -1.33 |
| ftsL | cell division protein FtsL | -1.29 |
| ribE | 6,7-dimethyl-8-ribityllumazine synthase | -1.29 |
| spa | staphylococcal protein A | -1.22 |
| ribB | 3,4-dihydroxy-2-butanone-4-phosphate synthase | -1.21 |
| lip1 | YSIRK domain-containing triacylglycerol lipase Lip1 | -1.13 |
| polX | DNA polymerase/3'-5' exonuclease PolX | -1.10 |
| secE | preprotein translocase subunit SecE | -1.07 |
| SAURSH1000_RS01225 | DUF5081 family protein | -6.66 |
| SAURSH1000_RS12995 | polysaccharide biosynthesis tyrosine autokinase | -6.40 |
| SAURSH1000_RS09780 | tRNA-Leu | -6.25 |
| SAURSH1000_RS07905 | hypothetical protein | -5.89 |
| SAURSH1000_RS11670 | hypothetical protein | -5.89 |
| SAURSH1000_RS07100 | hypothetical protein | -5.68 |
| SAURSH1000_RS00245 | tandem-type lipoprotein | -5.42 |
| SAURSH1000_RS04600 | YxeA family protein | -5.42 |
| SAURSH1000_RS12720 | hypothetical protein | -5.42 |
| SAURSH1000_RS02815 | hypothetical protein | -4.70 |
| SAURSH1000_RS06095 | hypothetical protein | -4.70 |
| SAURSH1000_RS10350 | 23S ribosomal RNA | -4.70 |
| SAURSH1000_RS08560 | DUF3969 family protein | -3.42 |
| SAURSH1000_RS00500 | replication initiation protein | -3.29 |
| SAURSH1000_RS09725 | ACT domain-containing protein | -2.97 |
| SAURSH1000_RS12020 | hypothetical protein | -2.81 |
| SAURSH1000_RS01625 | DUF951 domain-containing protein | -2.51 |

| | | |
|---|---|---|
| SAURSH1000_RS01815 | superantigen-like protein SSL3 | -2.40 |
| SAURSH1000_RS03425 | aminodeoxychorismate/anthranilate synthase component II | -2.22 |
| SAURSH1000_RS04605 | ABC transporter ATP-binding protein | -2.08 |
| SAURSH1000_RS12800 | hypothetical protein | -2.04 |
| SAURSH1000_RS01615 | ParB/RepB/Spo0J family partition protein | -1.99 |
| SAURSH1000_RS04955 | ABC transporter permease | -1.86 |
| SAURSH1000_RS00635 | ABC transporter substrate-binding protein | -1.74 |
| SAURSH1000_RS03555 | siderophore ABC transporter substrate-binding protein | -1.70 |
| SAURSH1000_RS09465 | membrane protein | -1.63 |
| SAURSH1000_RS04085 | teichoic acid D-Ala incorporation-associated protein DltX | -1.63 |
| SAURSH1000_RS11415 | N-acetyltransferase | -1.60 |
| SAURSH1000_RS00205 | flavodoxin family protein | -1.55 |
| SAURSH1000_RS00765 | RES domain-containing protein | -1.53 |
| SAURSH1000_RS02465 | Mini-ribonuclease 3 | -1.48 |
| SAURSH1000_RS08870 | helix-turn-helix transcriptional regulator | -1.48 |
| SAURSH1000_RS01855 | superantigen-like protein SSL10 | -1.46 |
| SAURSH1000_RS11705 | pyridoxal phosphate-dependent aminotransferase family protein | -1.46 |
| SAURSH1000_RS08405 | riboflavin synthase | -1.38 |
| SAURSH1000_RS02475 | NYN domain-containing protein | -1.30 |
| SAURSH1000_RS11390 | DUF3021 domain-containing protein | -1.26 |
| SAURSH1000_RS10620 | DUF6414 family protein | -1.25 |
| SAURSH1000_RS01620 | mechanosensitive ion channel family protein | -1.20 |
| SAURSH1000_RS02715 | DUF5327 family protein | -1.20 |
| SAURSH1000_RS08545 | excalibur calcium-binding domain-containing protein | -1.19 |
| SAURSH1000_RS00760 | sce7725 family protein | -1.16 |
| SAURSH1000_RS03575 | EMYY motif lipoprotein | -1.08 |
| SAURSH1000_RS06555 | 5-bromo-4-chloroindolyl phosphate hydrolysis family protein | -1.06 |

**Table S4.** DEGs of SWe-treated *S. aureus* compared to negative control at 24 h (*p* <0.05)

| Gene | Function | Log2FC |
|---|---|---|
| | | 24h |
| kdpF | K(+)-transporting ATPase subunit F | 4.83 |
| asp3 | accessory Sec system protein Asp3 | 3.28 |
| nrdF | class 1b ribonucleoside-diphosphate reductase subunit beta | 3.03 |
| gatC | Asp-tRNA(Asn)/Glu-tRNA(Gln) amidotransferase subunit GatC | 2.97 |
| yut | urea transporter | 2.79 |
| lacA | galactose-6-phosphate isomerase subunit LacA | 2.72 |
| nrdI | class Ib ribonucleoside-diphosphate reductase assembly flavoprotein NrdI | 2.40 |
| nrdE | class 1b ribonucleoside-diphosphate reductase subunit alpha | 2.39 |
| gcvpB | aminomethyl-transferring glycine dehydrogenase subunit GcvPB | 2.30 |
| gcvpA | aminomethyl-transferring glycine dehydrogenase subunit GcvPA | 2.29 |
| asp1 | accessory Sec system protein Asp1 | 2.19 |
| budA | acetolactate decarboxylase | 2.18 |
| asp2 | accessory Sec system protein Asp2 | 2.17 |
| coa | staphylocoagulase | 2.08 |
| rpmD | 50S ribosomal protein L30 | 2.06 |
| alsS | acetolactate synthase AlsS | 2.05 |
| essA | type VII secretion protein EssA | 1.98 |
| mnhe1 | Na+/H+ antiporter Mnh1 subunit E | 1.86 |
| rplR | 50S ribosomal protein L18 | 1.84 |
| thiD | bifunctional hydroxymethylpyrimidine kinase/phosphomethylpyrimidine kinase | 1.82 |
| xpt | xanthine phosphoribosyltransferase | 1.78 |
| mnhG1 | Na+/H+ antiporter Mnh1 subunit G | 1.77 |
| gpmI | 2,3-bisphosphoglycerate-independent phosphoglycerate mutase | 1.74 |
| putp | sodium/proline symporter PutP | 1.74 |
| atpD | F0F1 ATP synthase subunit beta | 1.73 |
| atpG | ATP synthase F1 subunit gamma | 1.73 |
| sasA | serine-rich repeat glycoprotein adhesin SasA | 1.71 |
| qoxC | cytochrome aa3 quinol oxidase subunit III | 1.70 |
| tpiA | triose-phosphate isomerase | 1.68 |
| SAURSH1000_RS00970 | hypothetical protein | 4.83 |
| SAURSH1000_RS10820 | hypothetical protein | 4.83 |
| SAURSH1000_RS00895 | hypothetical protein | 4.38 |
| SAURSH1000_RS02815 | hypothetical protein | 3.77 |
| SAURSH1000_RS09665 | YeeE/YedE family protein | 3.77 |
| SAURSH1000_RS00910 | acyl-CoA dehydrogenase family protein | 3.42 |
| SAURSH1000_RS07905 | hypothetical protein | 3.19 |
| SAURSH1000_RS00980 | PTS sugar transporter subunit IIA | 3.00 |
| SAURSH1000_RS01085 | 6-phospho-beta-glucosidase | 2.90 |
| SAURSH1000_RS01080 | glucose PTS transporter subunit IIA | 2.75 |
| SAURSH1000_RS12065 | hypothetical protein | 2.63 |
| SAURSH1000_RS00915 | class I adenylate-forming enzyme family protein | 2.63 |
| SAURSH1000_RS00655 | NAD-dependent formate dehydrogenase | 2.40 |
| SAURSH1000_RS00920 | acyl CoA:acetate/3-ketoacid CoA transferase | 2.28 |
| SAURSH1000_RS03020 | metal ABC transporter ATP-binding protein | 2.26 |
| SAURSH1000_RS09955 | single-stranded DNA-binding protein | 2.22 |
| SAURSH1000_RS04360 | ABC transporter permease | 2.22 |

| | | |
|---|---|---|
| SAURSH1000_RS12775 | hypothetical protein | 2.21 |
| SAURSH1000_RS04375 | peptide ABC transporter substrate-binding protein | 2.11 |
| SAURSH1000_RS00975 | BglG family transcription antiterminator | 2.11 |
| SAURSH1000_RS04840 | ECF transporter S component | 2.09 |
| SAURSH1000_RS02810 | DUF443 domain-containing protein | 2.06 |
| SAURSH1000_RS02715 | DUF5327 family protein | 2.05 |
| SAURSH1000_RS04370 | ATP-binding cassette domain-containing protein | 2.03 |
| SAURSH1000_RS12470 | acyltransferase family protein | 1.94 |
| SAURSH1000_RS04355 | ABC transporter permease | 1.92 |
| SAURSH1000_RS01465 | PTS ascorbate transporter subunit IIC | 1.89 |
| SAURSH1000_RS00995 | zinc-binding dehydrogenase | 1.88 |
| SAURSH1000_RS04365 | ABC transporter ATP-binding protein | 1.86 |
| SAURSH1000_RS01060 | response regulator transcription factor LytR | 1.79 |
| SAURSH1000_RS00990 | PTS galactitol transporter subunit IIC | 1.76 |
| SAURSH1000_RS11960 | AbgT family transporter | 1.72 |
| SAURSH1000_RS07570 | divalent metal cation transporter | 1.71 |
| SAURSH1000_RS03015 | metal ABC transporter permease | 1.68 |
| adhP | alcohol dehydrogenase AdhP | -2.76 |
| lukG | bi-component leukocidin LukGH subunit G | -2.72 |
| grpE | nucleotide exchange factor GrpE | -2.58 |
| nrdG | anaerobic ribonucleoside-triphosphate reductase activating protein | -2.51 |
| nrdD | anaerobic ribonucleoside-triphosphate reductase | -2.36 |
| argG | argininosuccinate synthase | -2.58 |
| argH | argininosuccinate lyase | -2.31 |
| clfB | MSCRAMM family adhesin clumping factor ClfB | -2.27 |
| rnpA | ribonuclease P protein component | -2.26 |
| clpB | ATP-dependent chaperone ClpB | -2.15 |
| purQ | phosphoribosylformylglycinamidine synthase I | -2.00 |
| prli42 | stressosome-associated protein Prli42 | -1.97 |
| groES | co-chaperone GroES | -1.97 |
| purE | 5-(carboxyamino)imidazole ribonucleotide mutase | -1.90 |
| uhpt | hexose-6-phosphate:phosphate antiporter | -1.89 |
| purK | 5-(carboxyamino)imidazole ribonucleotide synthase | -1.89 |
| clpP | ATP-dependent Clp endopeptidase proteolytic subunit ClpP | -1.85 |
| rimP | ribosome maturation factor RimP | -1.82 |
| cntK | histidine racemase CntK | -1.81 |
| mprF | bifunctional lysylphosphatidylglycerol flippase/synthetase MprF | -1.80 |
| yycF | response regulator YycF | -1.77 |
| pdxS | pyridoxal 5'-phosphate synthase lyase subunit PdxS | -1.77 |
| spxA | transcriptional regulator SpxA | -1.74 |
| brnQ | branched-chain amino acid transport system II carrier protein | -2.51 |
| hrcA | heat-inducible transcriptional repressor HrcA | -2.43 |
| isaB | immunodominant staphylococcal antigen IsaB | -2.36 |
| SAURSH1000_RS06565 | transposase | -6.02 |
| SAURSH1000_RS06030 | hypothetical protein | -5.45 |
| SAURSH1000_RS03655 | hypothetical protein | -5.19 |
| SAURSH1000_RS08580 | hypothetical protein | -5.19 |
| SAURSH1000_RS12350 | hypothetical protein | -4.92 |
| SAURSH1000_RS12855 | hypothetical protein | -4.88 |
| SAURSH1000_RS13280 | MAP domain-containing protein | -4.45 |

| | | |
|---|---|---|
| SAURSH1000_RS00255 | tandem-type lipoprotein | -4.14 |
| SAURSH1000_RS00885 | complement inhibitor SCIN family protein | -3.30 |
| SAURSH1000_RS01830 | superantigen-like protein SSL5 | -3.15 |
| SAURSH1000_RS06105 | hypothetical protein | -3.09 |
| SAURSH1000_RS04760 | hypothetical protein | -3.09 |
| SAURSH1000_RS12520 | DUF2316 family protein | -3.09 |
| SAURSH1000_RS12000 | hypothetical protein | -3.06 |
| SAURSH1000_RS05295 | superantigen-like protein SSL14 | -2.96 |
| SAURSH1000_RS02160 | GIY-YIG nuclease family protein | -2.82 |
| SAURSH1000_RS02695 | hypothetical protein | -2.75 |
| SAURSH1000_RS04625 | DoxX family protein | -2.66 |
| SAURSH1000_RS05330 | tRNA-Arg | -2.66 |
| SAURSH1000_RS06445 | hypothetical protein | -2.66 |
| SAURSH1000_RS08370 | MarR family transcriptional regulator | -2.61 |
| SAURSH1000_RS12175 | ATP-binding cassette domain-containing protein | -2.55 |
| SAURSH1000_RS02690 | protein VraC | -2.48 |
| SAURSH1000_RS01850 | superantigen-like protein SSL9 | -2.48 |
| SAURSH1000_RS08465 | transaldolase | -2.38 |
| SAURSH1000_RS10310 | hypothetical protein | -2.33 |
| SAURSH1000_RS12180 | ABC transporter permease | -2.27 |
| SAURSH1000_RS10170 | Dps family protein | -2.23 |
| SAURSH1000_RS07985 | hypothetical protein | -2.18 |
| SAURSH1000_RS03860 | cold-shock protein | -2.16 |
| SAURSH1000_RS05235 | hypothetical protein | -2.15 |
| SAURSH1000_RS04555 | competence protein ComK | -2.10 |
| SAURSH1000_RS08570 | ImmA/IrrE family metallo-endopeptidase | -1.97 |
| SAURSH1000_RS04880 | cytochrome ubiquinol oxidase subunit I | -1.95 |
| SAURSH1000_RS04885 | cytochrome d ubiquinol oxidase subunit II | -1.88 |
| SAURSH1000_RS04155 | FAD/NAD(P)-binding protein | -1.93 |
| SAURSH1000_RS06015 | MerR family transcriptional regulator | -1.88 |
| SAURSH1000_RS06585 | DUF6501 family protein | -1.87 |
| SAURSH1000_RS06590 | hypothetical protein | -1.86 |
| SAURSH1000_RS06350 | Y-family DNA polymerase | -1.82 |
| SAURSH1000_RS02420 | CtsR family transcriptional regulator | -1.81 |
| SAURSH1000_RS01980 | sodium-dependent transporter | -1.79 |
| SAURSH1000_RS11610 | type II toxin-antitoxin system Phd/YefM family antitoxin | -1.79 |
| SAURSH1000_RS00430 | hypothetical protein | -1.79 |
| SAURSH1000_RS02685 | thiolase family protein | -1.78 |
| SAURSH1000_RS11505 | nitrate/nitrite transporter | -1.73 |
| SAURSH1000_RS02670 | HAD family hydrolase | -1.72 |
| SAURSH1000_RS12830 | Crp/Fnr family transcriptional regulator | -1.72 |
| SAURSH1000_RS11605 | Txe/YoeB family addiction module toxin | -1.71 |
| SAURSH1000_RS02425 | UvrB/UvrC motif-containing protein | -1.70 |
| SAURSH1000_RS08255 | DUF948 domain-containing protein | -1.69 |
| SAURSH1000_RS04330 | MAP domain-containing protein | -1.68 |
| SAURSH1000_RS05320 | TDT family transporter | -1.67 |
| SAURSH1000_RS06180 | hypothetical protein | -1.66 |
